# SOX11 is a lineage-dependency factor and master epigenetic regulator in neuroblastoma

**DOI:** 10.1101/2020.08.21.261131

**Authors:** Bieke Decaesteker, Amber Louwagie, Siebe Loontiens, Fanny De Vloed, Juliette Roels, Suzanne Vanhauwaert, Sara De Brouwer, Ellen Sanders, Geertrui Denecker, Eva D’haene, Stéphane Van Haver, Wouter Van Loocke, Jo Van Dorpe, David Creytens, Nadine Van Roy, Tim Pieters, Christophe Van Neste, Matthias Fischer, Pieter Van Vlierberghe, Stephen S Roberts, Johannes Schulte, Sara Ek, Rogier Versteeg, Jan Koster, Johan van Nes, Katleen De Preter, Frank Speleman

## Abstract

The pediatric extra-cranial tumor neuroblastoma (NB) is characterised by a low mutation burden while copy number alterations are present in most high-risk cases. We identified SOX11 as a strong lineage dependency transcription factor in adrenergic NB based on recurrent chromosome 2p focal gains and amplifications, its specific expression in the normal sympatho-adrenal lineage and adrenergic NBs and its regulation by multiple adrenergic specific cis-interacting (super-)enhancers. Adrenergic NBs are strongly dependent on high *SOX11* expression levels for growth and proliferation. Through genome-wide DNA-binding and transcriptome analysis, we identified and validated functional SOX11 target genes, several of which implicated in chromatin remodeling and epigenetic modification. SOX11 controls chromatin accessibility predominantly affecting distal adrenergic lineage-specific enhancers marked by binding sites of the adrenergic core regulatory circuitry. During normal sympathoblast differentiation we find expression of SOX11 prior to members of the adrenergic core regulatory circuitry. Given the broad control of SOX11 of multiple epigenetic regulatory complexes and its presumed pioneer factor function, we propose that adrenergic NB cells have co-opted the normal role of SOX11 as a crucial regulator of chromatin accessibility and cell identity.

## INTRODUCTION

Neuroblastoma (NB) is the most common extra-cranial solid childhood cancer, originating from the developing sympatho-adrenergic nervous system^1^. The genomic landscape of NB is characterized by a low mutational burden and highly recurrent structural rearrangements. NB is considered a developmental disorder that is controlled by the complex interplay of multiple transcription factors (TFs) and reshaping of epigenetic landscapes^1^. Tumor cells can co-opt normal developmental pathways for functions that are linked to tumor progression and may become addicted to survival mechanisms controlled by developmental master regulators. Recent studies in NB revealed two distinct super-enhancer-associated differentiation states, i.e. adrenergic (ADRN) and early neural crest/mesenchymal (MES), each programmed by a specific core regulatory circuitry (CRC) defined by multiple lineage-specific transcription factors^2,3^. Furthermore, lineage identity switching and plasticity is an emerging key factor in therapy resistance of several cancers. Therefore, further insights into the nature and contribution of lineage dependency transcription factors may be important to understand frequently occurring relapses in high-risk NBs^4^.

Given that (1) in other tumors oncogenic lineage dependency factors were overexpressed through amplification^4^ and that (2) in addition to frequent MYCN amplification also other oncogenic (co-)drivers were found to be amplified in NB^1^, we aimed to identify novel putative lineage dependency TFs implicated in NB by delineating rare focal copy number gain and amplification events. We identified the SRY-related HMG-box transcription factor 11 (*SOX11*) as the sole protein coding gene residing in the shortest region of overlap at chromosome 2p distal to *MYCN* and pinpointed SOX11 as an important transcription factor implicated in adrenergic NB development.

SOX11 belongs to the SOX family of proteins, which are critical regulators of many developmental processes, including neurogenesis^5^. These TFs bind and bend the minor groove of the DNA using their highly similar high mobility group (HMG) domains. SOX11 belongs to the SoxC subgroup, which also includes SOX4 and SOX12, and the expression of these proteins is of key importance for the survival and development of the neural crest, multipotent neural and mesenchymal progenitors, and the sympathetic nervous system^6,7^. In addition to its presumed canonical transcription factor activity, SOX11 was recently also shown to have pioneering activity and thus can be assumed to direct chromatin accessibility at loci controlling cell fates^8^. Here, we report that *SOX11* acts as a lineage-dependency factor in adrenergic NB cells. We performed in depth analysis of functional SOX11 target genes which include multiple genes involved in chromatin remodeling and modification and DNA methylation and show that SOX11 impacts chromatin accessibility of lineage-specific adrenergic enhancers. Furthermore, a comparative analysis of SOX11 and MYCN binding sites suggests functional interaction between both transcription factors.

## RESULTS

### Rare focal amplifications and lineage-specific expression of *SOX11* in neuroblastoma

To identify lineage dependency TFs implicated in NB, we reanalysed DNA copy number profiling data of 556 high-risk primary NB tumors^9^ together with those from 263 additional published and 223 unpublished NB tumors^10,11^ and 39 neuroblastoma (NB) cell lines. We specifically searched for focal gains and/or amplifications of chromosomal segments encompassing TFs with a putative or known role in normal (neuronal) development. Within the 270 kb shortest region of overlap of the commonly gained large chromosome 2p segment (31% of cases), we identified focal amplifications (three primary NB cases with log2 ratio > 2) and high-level gains (two primary NB cases and one cell line with log2 ratio > 0.5) encompassing the TF *SOX11* as the only protein coding gene (Fig. 1a). Furthermore, all tumors showing *SOX11* focal amplification or high-level gain were also *MYCN* amplified (Fig. S1a). FISH analysis could be analyzed in two tumors and showed that *SOX11* and *MYCN* reside in two independently amplified segments (Fig. 1b). *SOX11* mRNA expression levels were found to be elevated in primary NB tumors with higher *SOX11* copy numbers (p-value = 1.82e-09, t-test) (n= 276) (Fig. 1c). Next, we observed that high *SOX11* mRNA expression levels (fourth quartile) are significantly related to worse overall and progression free survival in two independent NB cohorts of 276 and 498 patients (Fig. 1d, Fig. S1b). In addition, SOX11 immunohistochemical analysis using two independent SOX11 antibodies (Fig. S1c) showed that high SOX11 protein expression levels are associated with worse overall survival in a series of 56 NB tumors (Fig. 1e).

**Figure 1:**
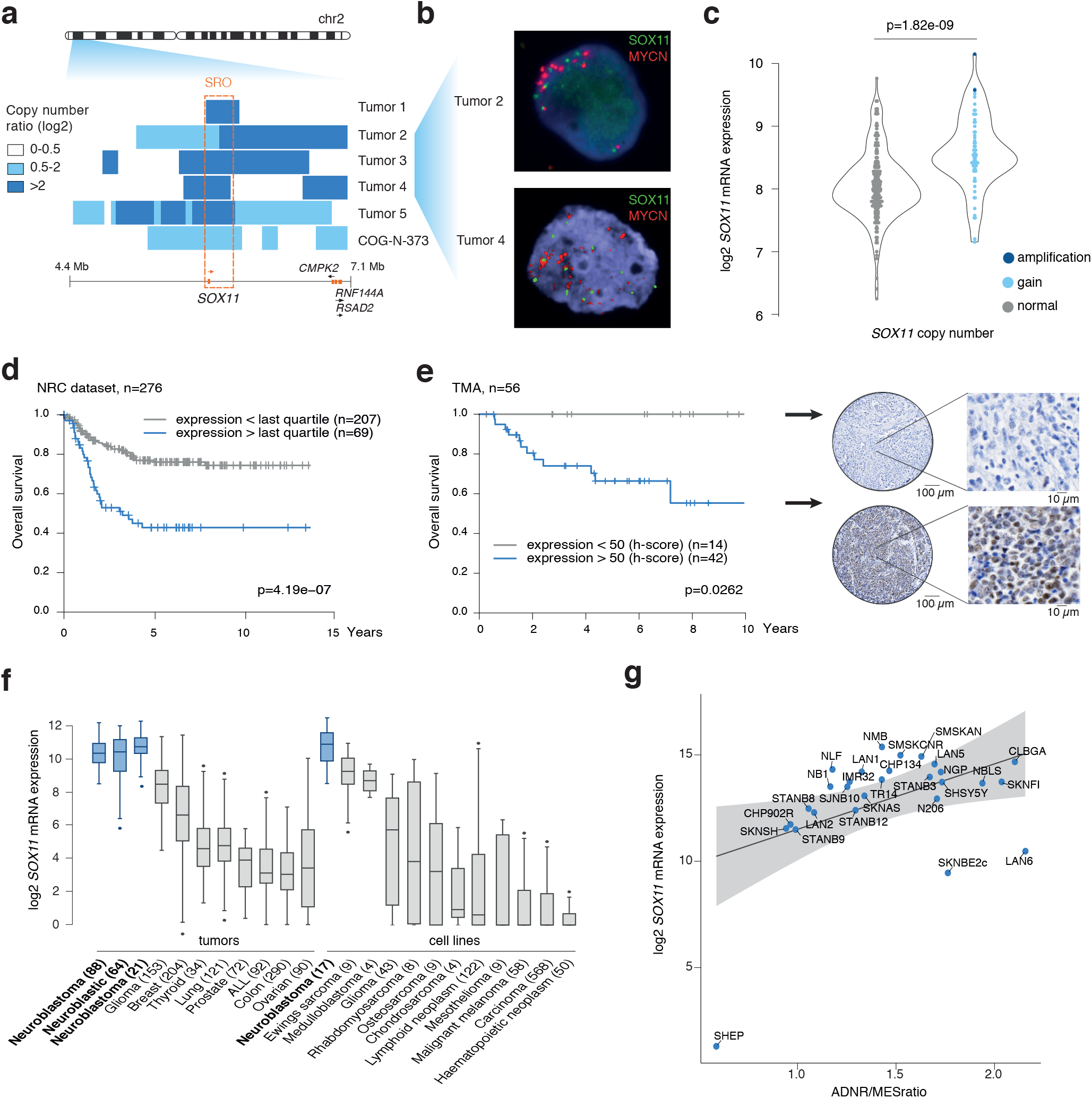
Rare focal amplifications and lineage-specific expression of *SOX11* in NB. A. Log2 copy number ratio on chromosome 2p (4.4–7.1Mb, hg19) showing shortest region of overlap (SRO) (chr2: 5,828,671-6,098,736; hg19) for high-level focal 2p gains (log2 ratio > 0.5) and amplification (log2 ratio > 2) in 5 NB tumors and 1 NB cell line encompassing the *SOX11* locus. Tumor 1 is evaluated by arrayCGH, tumor 5 by whole genome sequencing, and tumor 2, 3, 4 and the cell line by shallow whole genome sequencing. B. FISH analysis shows independent amplification of *SOX11* (green) and MYCN (red) in two of the tumor cases with *SOX11* amplification from Fig. 1A (asterix). C. *SOX11* (log2) mRNA levels according to copy number status (amplification (log2ratio>2), gain (log2ratio>0.3) or normal) in a cohort of 276 patients (NRC cohort, GSE85047) (T-test, p=1.82e-09). The two samples with *SOX11* amplification are tumor 4 and 5 from Fig. 1A. D. Kaplan-Meier analysis (overall survival) of 276 neuroblastoma patients (NRC NB tumor cohort, GSE85047) with high or low *SOX11* (log2) expression (using highest quartile as cut-off) (p=4.19e-07). E. Immunohistochemical staining for SOX11 on a tissue Micro-array (TMA) of 56 NB tumors and correlation of SOX11 protein levels (cut-off of H-score 50) with overall worse survival. For each group immunohistochemical staining of one tumor is depicted on the right (p=0.0262). F. *SOX11* expression (log2) in NB tumors and NB cell lines (blue) as compared to tumors or cell lines from other entities. Boxplots show the 1st quartile up to the 3rd quartile of the data values and median as a line within the box. Whiskers represent the values of the outer two quartiles maximized at 1.5 times the size of the box. If one or more values outside of the whiskers are present, then this is indicated with a single dot next to the implicated whisker. G. Log2 *SOX11* (log2) mRNA expression according to ADRN/MES ratio of the activity score (rankSum) for ADRN and MES signature^2^ in NB cell lines based on mRNA expression levels (n=28, p=3.94e-24, r=0.889, pearson).

*SOX11* is significantly higher expressed in high risk *MYCN* amplified tumors as compared to high risk *MYCN* non-amplified tumors and low risk tumors (Fig. S1d). Analysis of *SOX11* tissue specific expression patterns in R2 platform (http://r2amc.nl) and the Cancer Cell Line Encyclopedia (CCLE) showed the highest *SOX11* expression levels, highest copy number ratio and lowest methylation levels in NB tumors and cell lines as compared to other entities (Fig. 1f, Fig. S1e-f). Lineage restricted expression was evident from high expression levels in human fetal neuroblasts and in sympathetic neuronal lineages during early development as compared to normal cortex from the adrenal gland (Fig. S1g) and temporal increase of SOX11 expression was also noted in mouse NB tumor models in early hyperplastic lesions and full-blown tumors as compared to normal adrenal gland (Fig. S1h-i). Higher expression levels of *SOX11* both at mRNA and protein level were observed in adrenergic NB cell lines compared to mesenchymal NB cell lines and tumors (Fig. 1g, Fig. S1j-k). Taken together, we identified recurrent focal copy number alterations of the SOX11 locus in *MYCN* amplified tumors and adrenergic lineage-specific *SOX11* expression levels that are associated with poor prognosis in NB patients.

### *SOX11* is flanked by multiple *cis*-interacting distal adrenergic super-enhancers

Transcription factors implicated in defining cell lineage and identity are typically under the control of super-enhancers^12^. SOX11 was previously identified as a super-enhancer-associated TF in adrenergic NB cell lines^2^. To further map putative (super-) enhancers regulating *SOX11* expression, we investigated the *SOX11* locus and its neighbouring region for H3K27ac marks in 23 NB cell lines, a non-malignant neural crest cell line (hNCC) and the breast cancer cell line MCF-7 as non-embryonal control^3,13^. Distal to the SOX11 locus, we noted a large (1.1 Mb) region without protein coding genes (gene desert) which was marked by multiple H3K27ac peaks, indicative of the presence of multiple active enhancers, and in 7/23 NB cell lines we could identify a super-enhancer region (Fig. 2a, Fig. S2a). In accordance with the absence of *SOX11* expression in the breast cancer cell line MCF-7 cell line, the non-malignant neural crest cell line and the mesenchymal/neural crest like NB cell lines (GI-ME-N, SH-EP), H3K27ac peaks were absent in the gene desert, although in the latter two NB cell lines *SOX11* promoter H3K27ac peaks were seen. In further support of this cell identity specific H3K27ac pattern, we looked into H3K27ac data before and after NOTCH3-driven adrenergic-to-mesenchymal transition in NB cells^14^. As shown in Fig. S2b, a reduction in H3K27ac peak size was noted for the *SOX11* associated super-enhancer after NOTCH3 induction and thus mesenchymal transition. Interestingly, the *Sciatic Injury Induced LincRNA Upregulator Of SOX11* (*SILC1)* lncRNA is transcribed from the super-enhancer region and has been shown to play a critical role for induction of *SOX11* during neurite outgrowth and neuron regeneration^15^. This lncRNA is strongly expressed in adrenergic NB cells and highly correlated to *SOX11* expression levels in both NB cell lines (p=0.00806, R=0.773, Spearman, n=11) and tumors (p=3.01e-12, R=0.308, Spearman, n=498) (Fig. 2b, Fig. S2c). Also, in the normal developing sympathetic lineage and across a wide panel of cancer cell lines and adrenergic/mesenchymal isogenic cell line pairs^2^, we observed a similar pattern of *SILC1* expression as for *SOX11* (Fig. 2c, Fig. S2d-e). To investigate looping and physical contact of the cell type-specific enhancers with the promoter of *SOX11* and *SILC1*, we performed 4C-seq analysis for the *SOX11* and *SILC1* locus in the CLB-GA (adrenergic) and SH-EP (mesenchymal) NB cell lines and observed looping in this highly active region between the enhancer loci, the *SOX11* and *SILC1* promoter in the adrenergic cell line CLB-GA while this interaction was not detectable in the mesenchymal cell line SH-EP (Fig. 2d, Fig. S2f). In summary, multiple adrenergic specific enhancers are flanking the *SOX11* locus with putative roles in SOX11 regulation and SOX11 mediated cell identity, including the lncRNA SILC1.

**Figure 2:**
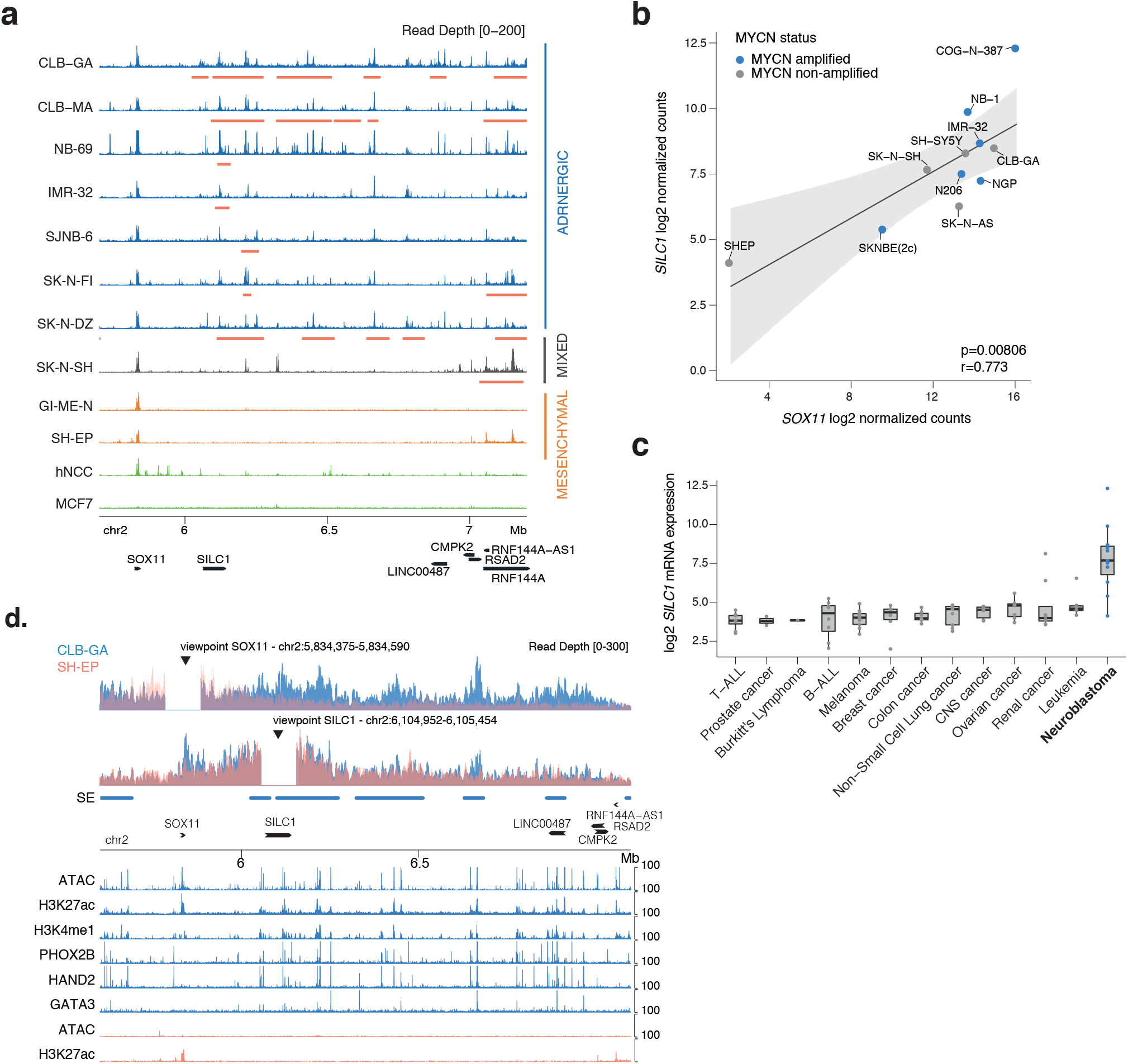
The *SOX11* locus is flanked by multiple adrenergic specific enhancers. A. H3K27ac activity in a region downstream of *SOX11* (chr2, 5.7–7.0Mb, hg19) in 6 adrenergic (CLB-GA, CLB-MA, NB-69, IMR-32, SJNB6, SKNFI, SKNDZ) and 1 mixed (SK-N-SH) (blue), 2 mesenchymal (SH-EP, GI-ME-N) (orange) and a non-malignant neural crest cell line hNCC and non-embryonal breast cancer cell line MCF-7 (green). Signal represents RPKM normalised ChIP signal, super-enhancers are annotated using LILY (red bar). B. *SOX11* mRNA correlation with *SILC1* mRNA levels in NB cell lines (n=11, p-value=0.00806, r=0.773, Spearman correlation) using the RNA Atlas (total RNA-sequencing). MYCN status is depicted for each cell line. C. *SILC1* (log2) mRNA expression in NB cell lines (blue) as compared to cell lines from other entities (gray). D. 4C-seq analysis of the promoter site and SE region downstream of SOX11 (chr2, 5.6-7.1Mb, hg19) in the NB cell lines CLB-GA (blue) and SH-EP (orange) with inclusion of published and unpublished ChIP tracks for H3K27ac, H3K4me1, PHOX2B, HAND2 and GATA3 in CLB-GA, and ATAC and H3K4me3 in SH-EP. Signal represents log likelihood ratio for the ChIP signal as compared to the input signal (RPM normalised). The viewpoint is located at the SOX11 TSS and SILC1 TSS site (cut out 100 kb). Super-enhancers of CLB-GA are annotated using LILY (blue bar).

### SOX11 is a lineage dependency factor in adrenergic NB cells

In a next step, we investigated whether adrenergic NB cells are dependent on *SOX11* expression for growth and survival as previously noted for MYCN and CRC members^16^. According to the publicly available CRISPR screen data in 769 cell lines (AVANA CRISPR 19Q1, available via the DepMap Portal)^17^, *SOX11* is identified as a strongly selective gene with dependency in 19 NB cell lines (p=6.6e-17) and more specifically in 13 *MYCN* amplified NB (p=2.3e-20) cell lines. We further assessed the phenotypic effects of *SOX11* knockdown in adrenergic NB cell lines, including two *MYCN* amplified cell lines (NGP and IMR-32) and two *MYCN* non-amplified cell lines with high MYC activity (SK-N-AS) or hTERT activation (CLB-GA), using RNA interference knockdown experiments (siRNAs and/or shRNAs) (Fig. 3a-b). Transient siRNA mediated knockdown of *SOX11* for 48h in NGP, SK-N-AS and CLB-GA cells resulted in the expected decreased number of colonies, as compared to transfected control cells (Fig. 3c). Concomitantly, long-term phenotype assessment after *SOX11* knockdown in the NGP, CLB-GA and IMR-32 cell lines, using the two most efficient shRNAs (sh3 and sh4), induced a significant G0/G1 growth arrest and reduction of proliferation (Fig. 3d-e). Our results confirm the CRISPR screen predicted strong lineage dependency role for SOX11 in the adrenergic NB cell line models we tested, both *MYCN* amplified and non-amplified.

**Figure 3:**
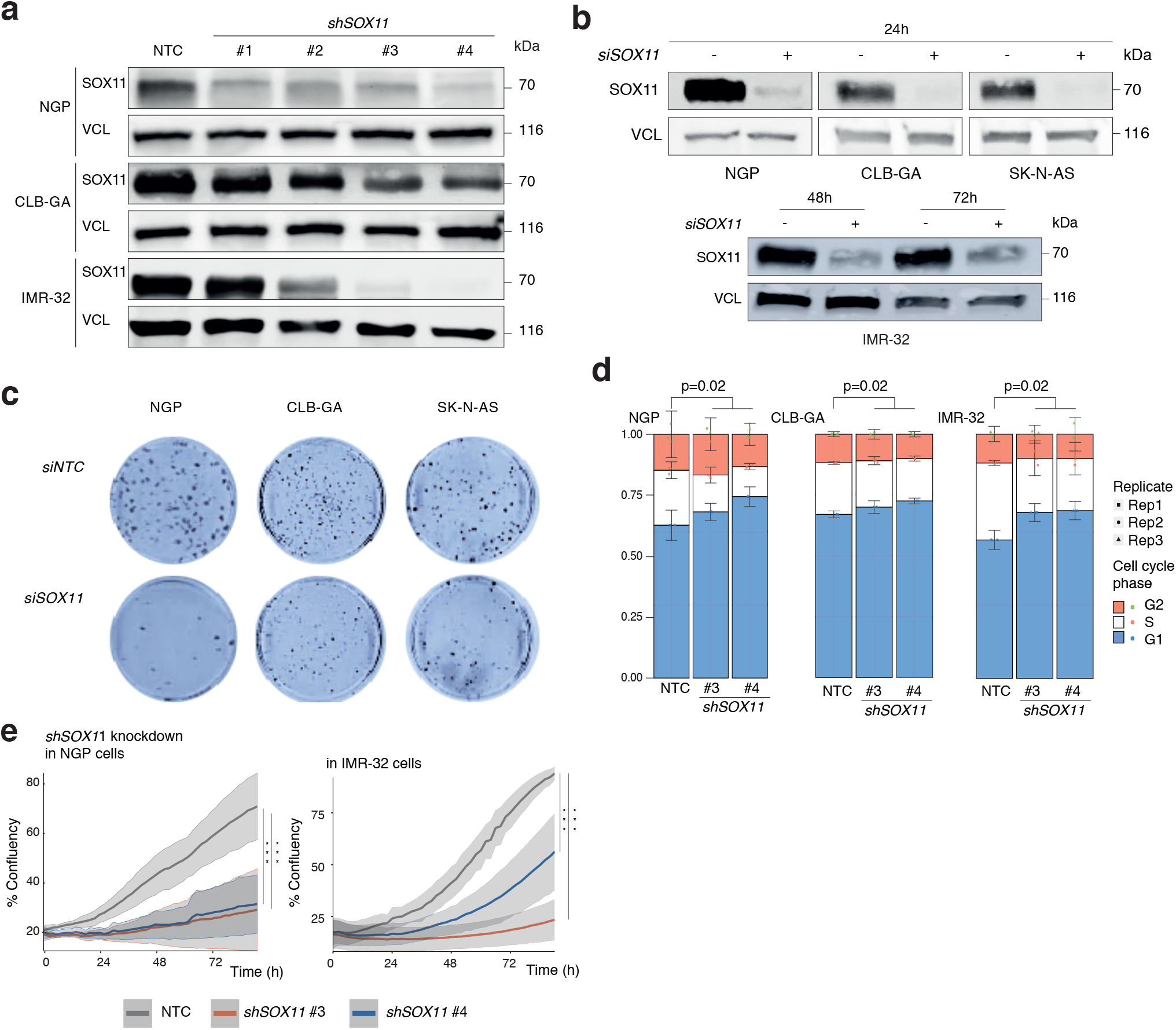
SOX11 is a lineage-dependency factor in adrenergic NB cells. A. SOX11 protein levels 6 days upon shSOX11 treatment in NGP, CLB-GA and IMR-32 cells with 4 different shRNAs and one non-targeting control (NTC). Vinculin (VCL) is used as loading control. B. SOX11 protein levels 48h and 72h upon si*SOX11* treatment in IMR-32 cells (nucleofection). SOX11 protein levels 24h upon siRNA treatment in NGP, CLB-GA and SK-N-AS cells (dharmafect transfection). Vinculin (VCL) is used as loading control. C. Reduction in colony formation capacity for NGP, CLB-GA and SK-N-AS cells, 14 days upon siRNA *SOX11* treatment (dharmafect transfection) as compared to non-targeting control (siNTC). D. Cell cycle analysis 6 days upon shRNA SOX11 treatment in the NGP and CLB-GA cell line. G1 cell cycle arrest upon *SOX11* knockdown as compared to the non-targeting control (NTC). Mann-Whitney statistical test is based on G1 phase percentage (shSOX11 vs NTC: NGP p-value=0.2, CLB-GA p-value=0.2, IMR-32 p-value=0.2). Data-points were mean-centered and auto scaled, error bars represent the 95% CI of three biological replicates for every cell line. E. Reduced proliferation (% confluence) over time upon prolonged *SOX11* knockdown with 2 different shRNAs for 4 days as compared to non-targeting control (NTC) in NGP and IMR-32 cells. Error bars represent the 95% CI of 3 biological replicates. ANOVA test followed by Tukey post-hoc test (NGP: sh3 vs NTC p=4.51e-14, sh4 vs NTC p=5.06e-14; IMR-32: sh3 vs NTC p = 2.16e-14, sh4 vs NTC p=2.74e-06).

### The SOX11 regulated transcriptome is involved in epigenetic control, cytoskeleton and neurodevelopment

To identify the key factors and pathways contributing to the adrenergic NB cell phenotype, we performed global transcriptome analysis upon transient siRNA-mediated knockdown of *SOX11* in IMR-32 cells for 48 hours (Fig. 4a, Fig. S4a, Supplementary Table 1). As an orthogonal experiment, we explored the transcriptome landscape after exogenously overexpressing *SOX11* for 48h in the mesenchymal SH-EP NB cell line (Fig. 4b, Fig. S4a, Supplementary Table 1). First, validating our *SOX11* knockdown specificity, we observed that both the significantly downregulated genes upon *SOX11* knockdown (n=1016, adj. pval <0.05) and the upregulated genes upon *SOX11* overexpression (n=4980, adj. pval <0.05) (further referred to as SOX11 activated targets) revealed enrichment of predicted SOX11 binding sites in their promoter (Fig. S4b). In line with the observed cell cycle arrest upon *SOX11* knockdown, the CDK inhibitor CDKN1A (also known as p21) (p=6.8e-07) was one of the top upregulated genes (Fig. S4c). Unexpectedly, multiple epigenetic modifiers involved in DNA and histone methylation, SWI/SNF and PRC1/2 complex members were strongly enriched, several of which were identified through ChIP sequencing as direct SOX11 targets (see further) (Fig. 4c, Fig. S4d). Of further interest and in concordance with the above mentioned putative epigenetic drivers regulated by SOX11, upregulated genes upon *SOX11* overexpression are involved in promoter and enhancer binding (Fig. 4c, Fig. S4e). In accordance with the putative role of SOX11 in adrenergic cell identity, we find enrichment for the genes of the proneuronal subtype in glioblastoma and adrenergic subtype in NB amongst the SOX11 activated targets. Vice versa, genes of the mesenchymal subtype are enriched amongst upregulated genes upon *SOX11* knockdown and downregulated genes upon SOX11 overexpression (further referred as the SOX11 repressed targets) (Fig. S4f)^18^. Altogether, this is suggestive of a possible role for SOX11 in maintenance of cell identity. Furthermore, gene set enrichment analysis for the SOX11 activated targets revealed strong enrichment for axon outgrowth, neural crest cell migration, cytoskeleton and RHO signalling (Fig. 4c, Fig. S4g). The strong induction of MARCKSL1 upon *SOX11* overexpression (log2 fold change 5.88) (Fig. S4c), an important regulator of actin stability and migration in neurons^19^, and the SOX11 regulated genes FSCN1 and TEAD2, two published SOX11 targets involved in cytoskeleton and neurodevelopment^7,20^, further supports the observed enriched gene sets.

**Figure 4:**
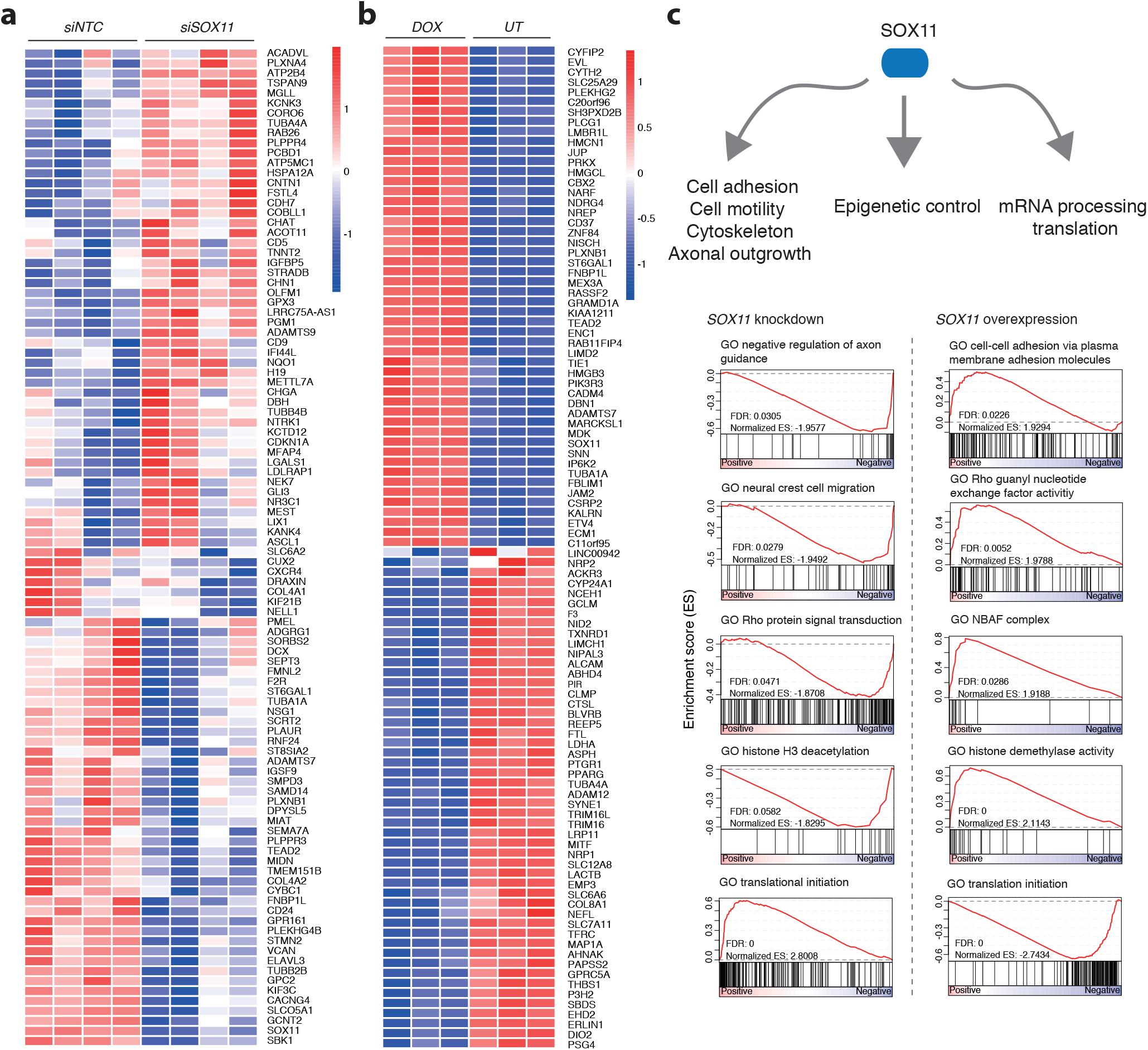
Cell motility, epigenetic control and translation are the hallmarks of the SOX11 regulated transcriptome. A. Top 50 up- and downregulated genes (based on limma voom t-value) upon SOX11 knockdown in IMR-32 cells represented in a heatmap. Heatmap color reflects row-wise z-score. B. Top 50 up- and downregulated genes (based on limma voom t-value) upon SOX11 overexpression in SH-EP cells represented in a heatmap. Heatmap color reflects row-wise z-score. C. Gene set enrichment analysis for the differentially expressed genes upon *SOX11* knockdown in IMR-32 cells and upon *SOX11* overexpression in SH-EP cells shows significant enrichment for genesets involved in cell adhesion, cytoskeleton, axonal outgrowth, epigenetic control, mRNA processing and translation.

Finally, SOX11 repressed targets are enriched for gene sets involved in translation initiation, ribosome hyperactivity and mRNA processing, which is indicative of an enhanced ribosome biogenesis response as previously reported in MYCN driven NB^21^(Fig. 4c). In conclusion, modulation of *SOX11* expression affects a broad range of phenotypic features in NB cells, including epigenetic control, cytoskeleton and neurodevelopment.

### SOX11 directly regulates the main modulators of the epigenome

To more comprehensively investigate direct functional targets of SOX11, we performed SOX11 DNA-occupancy analysis (ChIP-sequencing) in one of the prototypical *MYCN* amplified adrenergic cell lines, IMR-32. DNA binding motif analysis revealed a *de novo* SOX motif (24% of targets, bionomial test, p=1e-223) in the significant SOX11 bound sites (n=3105) providing support for the validity of our analytical procedure (Fig. S5a) (Supplementary Table 2). The majority of these SOX11 peaks showed significant overlap with the H3K27ac mark for active chromatin (88%), promoter mark H3K4me3 (67%) and open chromatin defined by ATAC-seq (95%)^13^, consistent with binding of SOX11 to both proximal and distal active transcriptional regulatory regions of both protein coding and noncoding genes (Fig. 5a-b, Fig. S5b) (p< 2.2e-16, Fisher exact test for overlap). We further validated the ChIP-seq targets by correlation with the above different generated transcriptome signatures upon *SOX11* knockdown and overexpression in a panel of 29 NB cell lines and in two primary NB tumor transcriptome datasets which showed strong overlap between SOX11 ChIP-seq activity score, the above transcriptome derived *SOX11* signatures, *SOX11* expression levels and NB patient survival outcome (Fig. 5c, Fig. S5c-e). Functional SOX11 targets were identified based on binding of SOX11 to their promotor/enhancer regions and when expression levels were significantly affected by SOX11 knockdown and overexpression. From a total of 2626 identified SOX11 binding sites, 313 functional targets were selected using these criteria. Strong enrichment of SOX11 binding was preferentially found in downregulated genes upon *SOX11* KD, suggesting that SOX11 acts mainly as transcriptional activator within this cell model (Fig. 5c-d). Also, further supporting the pathway analysis of the *SOX11* transcriptome data, the activated targets bound by SOX11 (n=234) showed chromatin remodeling, transcriptional regulation and axonogenesis as the most significantly enriched pathways (Fig. 5e, Fig. S5f). Next, we narrowed down the target list and established an 88-gene signature with SOX11 top direct functional targets containing genes with SOX11 DNA binding, differential expression upon SOX11 perturbation and positive correlation with *SOX11* expression in 2 independent primary NB cohorts (Fig. 5f, Supplementary Table 3). Strongly bound SOX11 targets included genes implicated in chromatin remodeling and enhancer activation (SWI/SNF core components SMARCC1, SMARCAD1), chromatin modification (PRC1/2-like complex components including CBX1 and CBX2), DNA methylation (DNMT1 binding partner UHRF1) and pioneer transcription factors including c-MYB, suggesting an important role for SOX11 in epigenetic regulation and control (Fig. 5b, Fig. 5g). In addition to c-MYB and SMARCC1, we observe elevated and reduced protein levels for the SWI/SNF core ATPase SMARCA4, upon *SOX11* overexpression and *SOX11* knockdown respectively (Fig. 5g, Fig. S5g-h). In summary, integration of SOX11 ChIP-seq data with SOX11 expression gene signatures reveals epigenetic modulators as bona-fide SOX11 direct targets in IMR-32.

**Figure 5:**
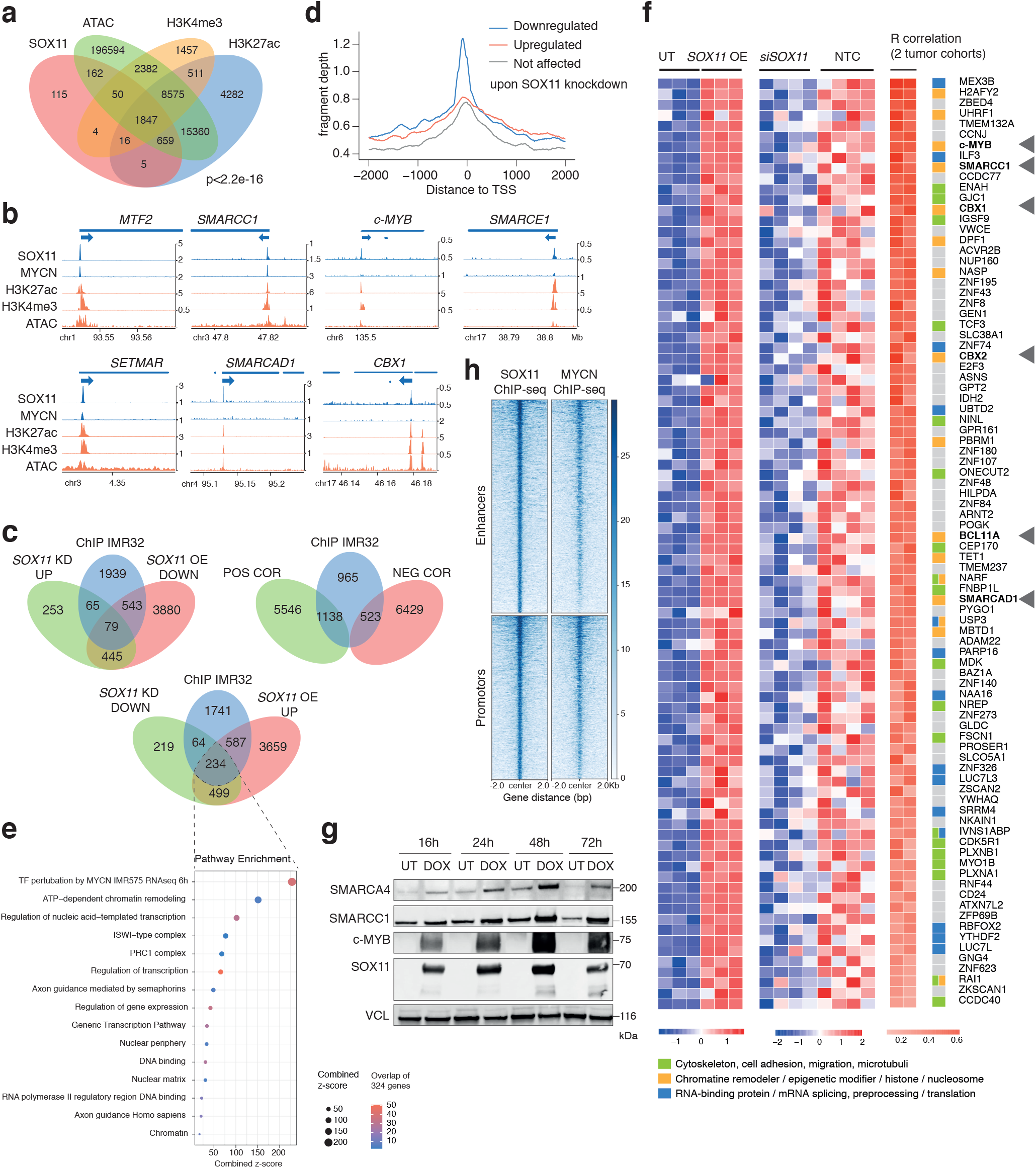
SOX11 is controlling the main modulators of the epigenome. A. Overlap (min. overlap = 20 bp) of the SOX11, H3K27ac, H3K4me3 ChIP-seq and ATAC peaks (MACS2 peakcalling qval < 0.05) in the IMR-32 cell line (Fisher test p-value < 2.2e^−16^). B. Binding of SOX11 and MYCN at *MTF2 (most significant target), SMARCC1, c-MYB, SMARCE1, SETMAR (2^nd^ highest significant target), SMARCAD1* and *CBX1* in IMR-32 cells and the overlap with H3K27ac, H3K4me3 and ATAC-seq peaks. Signal represents log likelihood ratio for the ChIP signal compared to input signal (RPM normalised). All peaks are called by MACS2 (q<0.05). C. Overlap of SOX11 ChIP-seq targets and genes perturbated upon *SOX11* overexpression and *SOX11* knockdown and overlap of the SOX11 ChIP-seq targets and the overlapping positively and negatively correlated genes in one or both NB tumor cohorts. Gene annotation for the SOX11 ChIP-seq targets is performed with Homer. D. Coverage of SOX11 ChIP signal for downregulated, upregulated genes and genes not affected (logFC) upon si*SOX11* knockdown in IMR-32 cells. Signal is depicted −2000 to +2000 bp distance from the peak summit. E. EnrichR analysis (http://amp.pharm.mssm.edu/Enrichr/) for SOX11 ChIP targets downregulated after SOX11 knockdown and upregulated after SOX11 overexpression. Depicted is the combined Z-score based on the multiplication of log p value computed with Fisher exact test and the z-score which is the deviation from the expected rank by the Fisher exact test (size), as well as the number of genes that overlap with the enriched genesets (color). F. 88 genes signature obtained by the overlap of upregulated genes upon *SOX11* knockdown, downregulated genes upon *SOX11* overexpression, SOX11 ChIP-seq targets, and positive correlation with *SOX11* expression in 2 different NB tumor cohorts (p-value < 0.05). The chromatin modifiers/remodelers CBX1, CBX2, SMARCC1, SMARCAD1, BCL11A and c-MYB are highlighted by an arrow. A color next to each gene represents the involved pathways. G. SMARCA4, SMARCC1, c-MYB and SOX11 and Vinculin protein levels in SH-EP cells upon *SOX11* overexpression. H. Heatmap profiles −2 kb and +2 kb around the summit of SOX11 ChIP-seq peaks. The heatmaps represent the ChIP-seq overlap of SOX11 and MYCN peaks in IMR-32, grouped for promoters or enhancers (homer annotation), and ranked according to the sums of the ChIP-seq peak scores across all ChIP-seq peaks in the heatmap.

### SOX11 acts in concert with MYCN to regulate a subset of downstream targets

The finding of a canonical MYCN motif (13.5% of the SOX11 targets, p=1e-28, Fig. S5a) and MAX motif (11.5% of the SOX11 targets, p=1e-32) in the SOX11 ChIP-seq targets (Supplementary Table 2). prompted us to perform ChIP-sequencing for MYCN in IMR-32 cells to explore the relation between SOX11 and MYCN in more depth. We observed an enrichment for both MYC(N) and SOX motifs in the identified MYCN targets in IMR-32 cells (Supplementary Table 2). Indeed, SOX11 and MYCN share common binding sites, both at promoters and enhancers (Fig. 5h) (Fisher exact test called peaks p= 5.5e-06). The enrichment for SOX11 bound activated genes among the genes downregulated 6h and 12h after inducible knockdown of *MYCN* in IMR-5/75 (Fig. S5f) further confirmed SOX11 and MYCN shared targets. Interestingly, SOX11-only bound genes at enhancers revealed strong enrichment for a non-canonical E-box motif, while SOX11-MYCN co-bound genes or MYCN-only bound genes at enhancers show enrichment for a canonical E-box motif (Supplementary Table 2). This suggests that SOX11 co-binds enhancers with bHLH TFs binding at non-canonical E-box motifs, such as the TFs *HAND2*, *TWIST1*, *TCF3* or *ASCL1* (Fig. S5a). Of further interest, genes that were directly bound by SOX11-MYCN showed enrichment for the genes downregulated 6h and 12h upon inducible knockdown of *MYCN* in IMR-5/75, as well as for gene sets involved in histone mediated repression, such as targets of EZH2, a common binding target of SOX11 and MYCN, and the SWI/SNF complex (Fig. S5i). In summary, our data indicates co-binding of SOX11 and MYCN which indicates a putative cooperative function between both genes.

### SOX11 regulates chromatin accessibility at active enhancers

SOX TFs exhibit the exceptional feature to bind the minor groove of DNA and subsequently bend the DNA, which is essential for the enhanceosome architecture as well as for the distortion of DNA wrapped around histones suggesting putative pioneer factor function^8^. As our current findings also reveal a role for SOX11 in transcriptional regulation of crucial components of several epigenetic protein regulatory complexes, including PRC2 and SWI/SNF, we propose a master epigenetic regulator function of SOX11 in adrenergic NB cells. To explore this further, we mapped chromatin accessibility changes by ATAC-sequencing 48h after *SOX11* knockdown in adrenergic CLB-GA cells. Differential ATAC-seq peaks in CLB-GA revealed only significant closed regions upon *SOX11* knockdown (n=2875), indicating a broad effect of SOX11 on chromatin accessibility. Of further interest, these sites of chromatin accessibility changes are predominantly observed at enhancers (Fig. 6a). Moreover, differential ATAC peaks overlapping with active chromatin marks (H3K27ac and H3K4me1 binding, n=1345) were highly enriched for motifs of the adrenergic core regulatory circuitry (CRC), including GATA3, PHOX2B, ISL1 and HAND2 (Fig. 6a). Indeed, 89% of the active enhancers closed upon *SOX11* knockdown overlap with binding of at least one member of the adrenergic CRC (PHOX2B, GATA3, HAND2) (Fig. S6a). Adrenergic specific super-enhancer regions^2^ show impaired chromatin accessibility upon *SOX11* knockdown (Fig. 6b). Given the adrenergic specific expression of SOX11, the presence of an active downstream super-enhancer, and the enrichment for CRC binding in the regions closed upon *SOX11* knockdown in adrenergic cells, we investigated the putative role of SOX11 in the adrenergic core regulatory circuitry. While the broad SOX11 enhancer binding activity also includes adrenergic CRC super-enhancers with lost chromatin accessibility upon *SOX11* knockdown (Fig. S6b), SOX11 binding is not enriched in enhancer regions that are occupied by several CRC members (Fig. S6c-d). In addition, SOX11 does not bind the super-enhancer and SILC1 lncRNA locus that loops to the SOX11 promotor. However, SOX11 binding is shown to occur at a more distal super-enhancer present in only a limited number of NB cell lines (3/23) (data not shown). Furthermore, we do not see transcriptional impact upon *SOX11* knockdown on the published CRC signatures or the key CRC members such as *GATA3* and *HAND2* (data not shown). To further understand the dynamic relation of SOX11 towards the CRC, we looked into expression levels of SOX11 and key CRC members PHOX2, HAND2 and GATA3 in human Pluripotent Stem Cells (hPSC) induced developing sympathoblasts (*Van haver et al., in preparation*). SOX11 was found to be expressed in earlier developmental stages (Fig 6c, Fig. S6e). Taken together, our findings support the notion that SOX11 is not a canonical CRC member but plays a distinct role during early sympathoblast development prior to emergence of the adrenergic master regulator PHOX2B and the other CRC members including HAND2 and GATA3. As SOX11 mediates chromatin accessibility for adrenergic core regulatory circuitry gene binding, we propose a role for SOX11 in the epigenetic establishment and/or maintenance of the adrenergic CRC through activation of various key epigenetic components.

**Figure 6:**
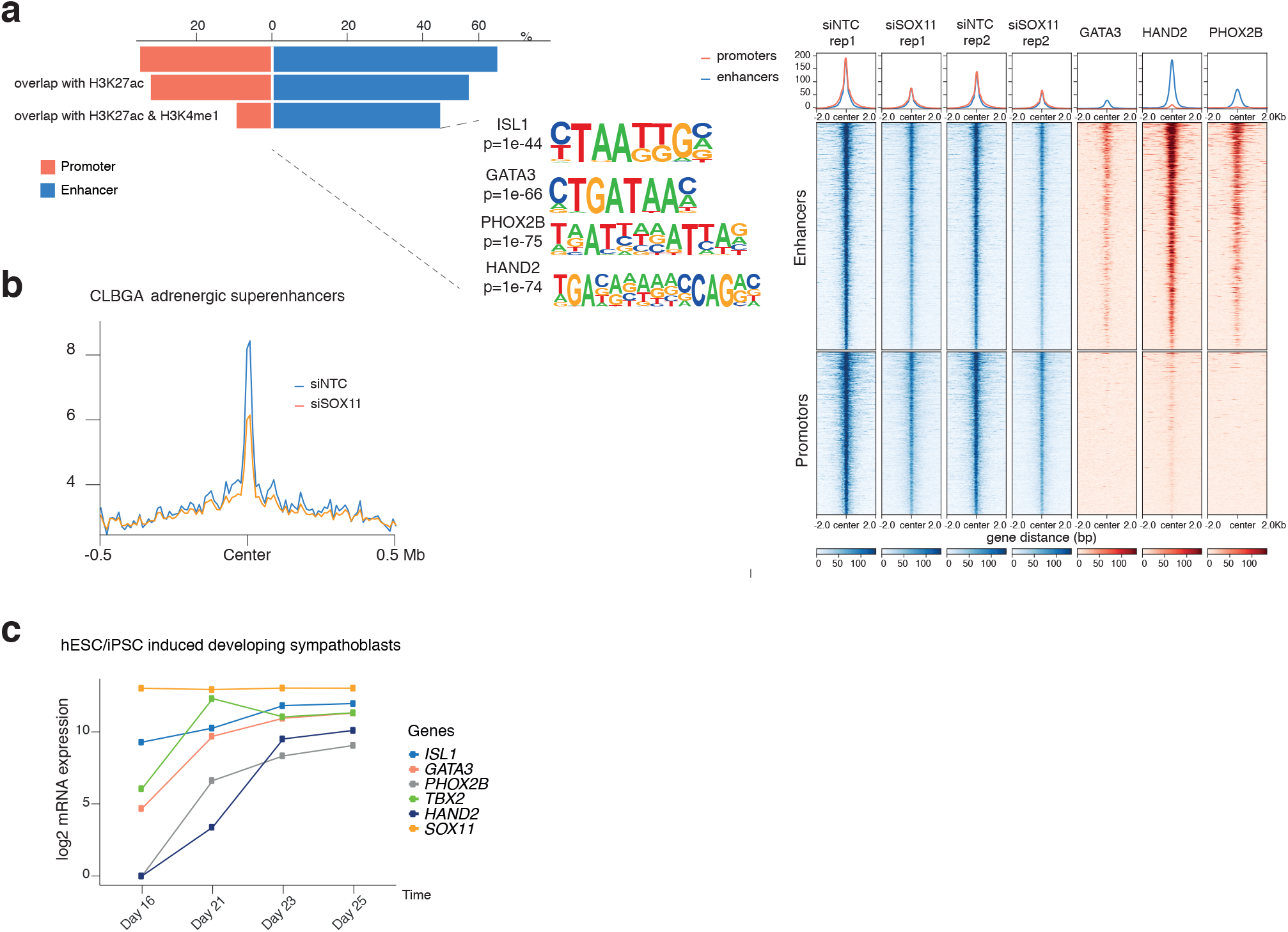
SOX11-mediated chromatin remodeling of lineage specific enhancers. A. ATAC-seq peaks with lost chromatin accessibility upon SOX11 knockdown in CLB-GA cells are annotated by homer as “promoter” or “enhancer”. For both classes overlap with H3K27ac and H3K4me1 is depicted. Motif enrichment of the CRC transcription factors PHOX2B, HAND2, ISL1 and GATA3 is found in the enhancers with lost chromatin accessibility overlapping with H3K27ac and H3K4me1. Heatmap profiles −2 kb and +2 kb around the summit of differential ATAC-seq peaks at enhancers or promoters upon SOX11 knockdown, depicting ATAC peaks with and without SOX11 knockdown (2 biological replicates), and GATA3, HAND2 and PHOX2B ChIP-seq profiles, ranked according to the sums of the peak scores across all peaks in the heatmap. Density profiles are shown representing the average ATAC- or ChIP-seq signal at the presented regions for promotors (orange) and enhancers (blue). B. Average RPKM normalized ATAC-seq reads upon siSOX11 knockdown (siSOX11, orange) as compared to non-targeting control (siNTC, blue) in CLB-GA cells, −0.5Mb and +0.5Mb around the center of adrenergic specific super-enhancers^2^. C. ISL1, GATA3, PHOX2B, TBX2, HAND2 and SOX11 (log2) expression during induced differentiation of hPSC cells along the sympatho-adrenal lineage. Expression levels depicted starting from day 16, upon sorting the cells for *SOX10* expression indicating cells committed to truncal neural crest cells, and followed-up during sympatho-adrenal development until day 25

## Discussion

Cellular mechanisms that govern lineage-specific proliferation and survival during development may be co-opted by tumor cells. Consequently, these tumors will be selectively dependent on such lineage factors offering interesting options for targeting these tumor cells whilst sparing normal tissues. We identified SOX11 as a lineage dependency gene in neuroblastoma (NB) which is recurrently affected by large segmental 2p gains in high risk NBs as well as recurrent focal gains and amplifications. SOX11 was identified as the sole protein coding gene residing in the shortest region of overlap at 2p distal to *MYCN*, suggesting a role as driver for selection of the respective amplicons during tumor formation. SOX11 is known to exhibit lineage specific expression during normal development, predominantly in the neuronal lineage. In line with the data from the CRISPR screen available through the DepMap portal, our *SOX11* knockdown data support that SOX11 is a dependency factor in NB. Furthermore, higher *SOX11* expression levels were found to be correlated with poor prognosis for NB patients. To gain insight into the functional contribution to the tumor phenotype, we identified functional SOX11 target genes through genome wide DNA binding combined with transcriptome analysis after *SOX11* knockdown and overexpression yielding 79 upregulated and 234 downregulated direct target genes indicating a primarily gene transcription activating role for SOX11. Both the diverse functionality of target genes and our *in vitro* experiments suggest a possible broad functional impact of SOX11 on the NB phenotype. Here, we further focussed on the possible role of SOX11 as epigenetic master regulator, given its direct role in regulation of components of the PRC2 and SWI/SNF complexes, the DNMT1 recruiting protein UHRF1 and pioneering TF c-MYB^22^. While these targets require further individual functional validation, the finding of multiple functional targets implicated in a broad range of essential epigenetic regulatory processes is intriguing. SOX11 was previously found to be affected by loss-of-function germline mutations causing the intellectual disability Coffin-Siris syndrome^23^, a disease mostly caused by mutations in SWI/SNF components including SMARCE1, and thus suggesting that SWI/SNF regulation is one of the major functions of SOX11 during neuronal development. We therefore propose that this function is also critically important in adrenergic NB cells. Possibly, SOX11 may contribute to the differentiation arrest in NB initiation as it has been reported that in normal development (e.g. cardiogenesis), activation of specific SWI/SNF components can have an impact on developmental transcriptional programs^24^. In this respect, it is of interest that overexpression of SMARCA4/BRG1, the catalytical component of the SWI/SNF complex, is essential for NB cell viability^25^. Furthermore, the previously established role of SWI/SNF chromatin remodeling in maintenance of lineage-specific enhancers leads us to hypothesize that the SWI/SNF complex could impact on NB tumor maintenance^26^.

While we observed an enrichment of adrenergic and mesenchymal gene signatures upon *SOX11* overexpression and knockdown respectively, SOX11 overexpression in itself was not sufficient to induce a transition in cell lineage, at least not at 48 hours after induction of SOX11 overexpression. The exact role of SOX11 in the development of the sympathetic neuronal lineage and its functional interconnection with the adrenergic TFs of the CRC will require further investigation. Our current data do not support an unequivocal role for SOX11 as a canonical CRC member, despite high *SOX11* expression being restricted to the adrenergic lineage. There is further no indication for SOX11 being part of the extended regulatory network (ERN), reported as downstream genes regulated by super-enhancers and the CRC transcriptional factors^27^, that do not necessarily bind to the super-enhancers of the CRC genes, nor have an autoregulatory function. Rather than a CRC TF like function, we propose a CRC upstream hierarchical function based on gene expression analysis of SOX10 positive neural crest derived maturing sympathetic adrenergic neuroblasts. While the adrenergic master regulator PHOX2B is strongly induced at day 23 of differentiation together with several CRC TFs including HAND2 and GATA3, SOX11 is clearly expressed much earlier from day 16 on of the differentiation track. Given the recently identified role for SOX11 as pioneer factor and recent insights into the interaction of non-pioneering factor to control to modulate cell fate and identity^28^, together with the potential of SOX11 to broadly activate multiple epigenetic regulatory protein complexes, we propose a critical direct or cooperative role for SOX11 in induction of the CRC of TFs in developing sympathoblasts. We also propose that several SOX11 driven functions are co-opted by the transformed neuroblasts contributing to the aggressive phenotype of high-risk adrenergic NBs. Further support for this view comes from the recent description of a unique aggressive transitional cell state important for the inter-transition between adrenergic and mesenchymal cells through single-cell RNA-sequencing analysis of peripheral neuroblastic tumors^29^. Interestingly, *SOX11* is described as a marker gene of the transitional state, amongst *MYCN* and others, while *GATA3* and *HAND2* mark the adrenergic state. This is in line with the fact that SOX11 was recently reported to be activated in a so-called proliferative active bridging population (transient cellular state) connecting a progenitor cell type coined Schwann cell precursors and their differentiated counterpart, chromaffin cells^30^, while the transitional signature in the above described tumors is enriched in the bridging population.

Our study also highlights long noncoding RNA *SILC1* as a putative regulator of SOX11 activity. SILC is marked by strong adrenergic-specific H3K27ac peaks and maps to a putative regulatory SOX11 distal protein coding poor region. Interestingly, the *SILC1* sequence and expression, in contrast to most other lncRNAs, is conserved in various mammals, and has been shown to regulate *SOX11* expression in *cis* during neurite outgrowth and neuron regeneration^15^. Additionally, this lncRNA was previously shown to be implicated in neuronal cells with evidence for chromatin looping towards the SOX11 promotor as shown by HiC genome wide chromatin conformation data in mice.^31,32^ Recent data also associated *SILC1* expression with proliferation and apoptosis in NB.^33^

In conclusion, we identify SOX11 as a novel dependency factor in adrenergic high-risk NB with a putative function as epigenetic master regulator upstream of the recently discovered CRC. Further studies in *in vitro* cellular models and targeted overexpression to the sympathetic adrenergic lineage in mice or zebrafish as well as developmental studies in animals or hPSC differentiation models are needed to further explore the complex interplay of the broad range of transcription factors in adrenergic neuroblasts and how these interact with unscheduled MYCN overexpression leading to NB formation.

## Material & Methods

### Samples and cell lines

All patient specimens and samples were used in accordance with institutional and national policies at the respective locations, with appropriate approval provided by the relevant ethical committees at the respective institutions. All patient-related information was anonymized.

Copy number analysis was performed on primary untreated NB tumors, representative of all genomic subtypes including 263 and 223 samples^10,11^ of the NRC cohort (GSE85047), 556 samples of the NB high-risk cohort^9^ (GSE103123), and one unpublished in-house sample. In addition, copy number data of 33 NB cell lines^10^ and the cell line COG-N-373h (Fig. 1a) were used. *SOX11* expression analysis was performed on 283 NB tumors for which copy number (n=218), mRNA expression (n=283) and patient survival (n=276) data were available from the Neuroblastoma Research Consortium (NRC, GSE85047), which is a collaboration between several European NB research groups. Additionally, the NB dataset from Su et al. (n=489, GSE45547) was used as validation cohort^34^.

All NB cell lines used in this manuscript (genotype and mutation status in Supplementary Table 4), were grown in RPMI1640 medium supplemented with 10% foetal bovine serum (FBS), 2 mM L-Glutamine and 100 IU/ml penicillin/streptavidin (referred further to as complete medium) at 37°C in a 5% CO2 humid atmosphere. Short Tandem Repeat (STR) genotyping was used to validate cell line authenticity prior to performing the described experiments and Mycoplasma testing was done every two months and no mycoplasma was detected.

### High-resolution DNA copy number analysis

DNA was obtained using the QiaAmp DNA Mini kit (Qiagen #51304) according to the manufacturer’s instructions and concentration was determined by Nanodrop (Thermo Scientific) measurement. Array comparative genomic hybridisation (arrayCGH) was performed as previously described^35^, as well as shallow whole genome sequencing^36^. Copy number data were processed, analysed and visualised using VIVAR^35^. Fluorescence in situ hybridization (FISH) was performed as previously described^37^ using the CTD-2037E22 probe for the *SOX11* locus. *SOX11* amplifications and high-level focal gains were identified as copy number segments overlapping with the *SOX11* locus with log2 ratio >= 2 and >= 0.3 respectively and a maximal size of 5 Mb.

### Tissue micro-array

A NB tissue micro-array was used as previously described^38^. For immunohistochemical staining, 5-μm sections were made, antigen retrieval was done in citrate buffer and endogenous peroxidases were blocked with H_2_O_2_ (DAKO). The sections were incubated with primary antibodies (SOX11-C1 antibody^39^ = Antibody 1, SOX11 antibody from Klinipath (cat#ILM3823-C01) = Antibody 2), followed by incubation with the Dako REALTM EnvisionTM-HRP Rabbit/Mouse system and substrate development was done with DAB (DAKO). Scanning of the slides was done using the Zeiss Axio Scan.Z1 (Zeiss) and counting of SOX11 positive NB cells was done by H-scoring. In brief, the percentage of SOX11-positive cells is each time multiplied by the intensity (0, 1, 2 of 3): [1 × (% cells 1+) + 2 × (% cells 2+) + 3 × (% cells 3+)]. Blind scoring was done by two independent persons. Each sample was present in triplicate and scores are presented as the average of the three replicates. 15 samples were omitted due to lack of survival data.

### Statistical and transcriptomic analysis of NB cohorts and other entities

Neuroblastoma transcriptomic analysis was performed on a dataset of 283 NB tumors for which copy number (n=275), mRNA expression (n=283) and survival (n=276) data were available from the Neuroblastoma Research Consortium (NRC, GSE85047), which is a collaboration between several European NB research groups. Additionally, the NB dataset from Su et al. (n=489, GSE45547) was used as validation cohort^34^. The *SILC1* expression data were analysed using the RNA Atlas (Lorenzi et al., in preparation). H3K27ac ChIP-seq data and super-enhancer annotation were public available from Boeva et al. ^3^ and Decaesteker et al. ^13^. The Depmap array^17^ and R2 platform (http://r2amc.nl) were used as repositories for gene expression and dependency data of different tumor entities.

All statistical analyses (two-sided t-test, Wilcoxon test, kruskal-wallis, ANOVA, post-hoc dun-test or tukey test, Kaplan-Meier, correlation spearman and pearson) were done using R (version 3.5.1). For correlation analysis, genes were ranked according to Pearson correlation coefficient.

### 4C-sequencing

4C-sequencing was performed as previously described^13^. For visualization purposes, the viewpoint was removed (chr2:58210000-4835000) and the plot was generated using R package Sushiplot with normalized bedgraph files.

### Transfection and nucleofection of cell cultures

Cells were seeded in 6-well tissue culture plates 24 hours prior to transfection. 100nM of siRNA non-targeting control (siRNA NTC, D-001810-10-05) or siRNA SOX11 (L-017377-01-0005, Dharmacon) were transiently transfected using DharmaFect 2 (Thermo Fisher Scientific) according to the manufacturer's guidelines. For nucleofection, cells were nucleofected with 100 nM of the above described siRNA NTC and siRNA SOX11 using the Neon Transfection System (Thermo Fisher Scientific) and subsequently seeded in 6-well or T25 tissue culture plates.

### Generation of stable cell lines

Four different mission shRNAs from the TRC1 library (Sigma-Aldrich, TRCN0000019174, TRCN0000019176, TRCN0000019177, TRCN0000019178, referred in the manuscript as sh1, sh2, sh3, sh4 respectively) targeting SOX11 and one non-targeting shRNA control (SHC002, NTC) were used to generate neuroblastoma cell lines with *SOX11* knockdown.

Virus was produced by seeding 3×10^6^ HEK-293TN cells in a 15cm^2^ dish 24h prior to transfection. Transfection of the cells was done with trans-lentiviral packaging mix and lentiviral transfection vector DNA according to the Thermo Scientific Trans-Lentiviral Packaging Kit (TLP5913) using CaCl_2_ and 2x HBSS. 16 hours after transfection, cells were refreshed with reduced serum medium and lentivirus-containing medium was harvested 48 hours later. Virus was concentrated by adding 2500 *μ*l ice-cold PEG-IT (System Biosciences) to 10 ml harvested supernatants and incubating overnight at 4°C, after which complete medium was added to the remaining pellet upon centrifugation. NGP, CLB-GA and IMR-32 cells were transduced by adding 250 *μ*l concentrated virus to 1750 *μ*l complete medium. 24h after transduction cells were refreshed with medium and 48h after transduction, cells were selected using 1 *μ*g/ml puromycin.

For SOX11 inducible overexpression, the OriGene vector SC303275 containing the cDNA of SOX11 was amplified by PCR and the obtained fragment was gel purified and ligated into the opened NdeI site of response vector pLVX-TRE2G-Zsgreen1 (Takara, cat#631353) producing pLVX-TRE3G-Zsgreen1-IRES-hSOX11. The constructed plasmid was verified by restriction digest and sequenced by Sanger DNA sequencing (GATC). Lenti-X 293T Cells (Takara, cat#632180) were transfected with the regulator vector pLVX-pEF1a-Tet3G (cat#631353 and Lenti-X Packaging Single Shots (VSV-G) (cat#631275) according to the manufacturer’s instructions. The supernatant containing the lenti-virus was collected, filtered through a 0.45 *μ*m filter and concentrated using PEG-IT. SH-EP cells were infected with the concentrated virus and 48 hours of incubation thereafter, the transduced cells were selected using 500 *μ*g/ml G418. Three individual clones were obtained by limiting dilution. After clonal expansion, the TET protein expression in each clone was checked by immunoblotting using TetR monoclonal antibody (Clone 9G9) (Clontech, cat#631131). In addition, induction of each expressing clone was tested after transduction with the pLVX-TRE3G-Luc control vector. Selected clones were transduced with lentivirus produced as described above from vector pLVX-TRE3G-Zsgreen1-IRES-hSOX11 and subsequently selected with 1 *μ*g/ml puromycin. The SH-EP SOX11 clones were grown in completed medium supplemented with 10% tetracyclin-free FBS to avoid leakage.

### Phenotypic assessment of cells

For the colony formation assay, 2000 viable NGP, CLB-GA and SK-N-AS cells with or without *SOX11* knockdown were seeded in a 6-cm dish in a total volume of 5 ml complete medium and were then left unaffected for 10-14 days at 37°C. After an initial evaluation under the microscope, the colonies were stained with 0.005% crystal violet and digitally counted using ImageJ. The IncuCyte^®^ Live Cell imaging system (Essen BioScience) was used for assessment of proliferation after *SOX11* knockdown or overexpression. Briefly, 15×10^3^ viable NGP or 10×10^3^ viable SH-EP cells, with or without *SOX11* knockdown or overexpression, were seeded in 5 replicates in a 96-well plate (Corning costar 3596) containing complete medium. Cell viability was measured in real-time using the IncuCyte by taking photos every 3 hours of the whole well (4x). Masking was done using the IncuCyte^®^ ZOOM Software.

For cell cycle analysis, 7×10^5^ cells were seeded in a T25 in complete medium and nucleofected with SOX11 siRNA or transduced with *SOX11* shRNAs and respectively controls and selected with puromycin, as described above. Cells were trypsinized and washed with PBS. The cells were resuspended in 300 *μ*l cold PBS and while vortexing, 700 *μ*l of 70% ice-cold ethanol was added dropwise to fix the cells. Following incubation of the sample for minimum 1 hour at −20 °C, cells were washed in PBS and resuspended in 500 *μ*l PBS with RNase A to a final concentration of 0.25 mg/ml. Upon 1 hour incubation at 37° C, 20 *μ*l Propidium Iodide solution was added to a final concentration of 40 *μ*g/ml. Samples were loaded on a BioRad S3^TM^ Cell sorter and analysed with the Dean-Jett-Fox algorithm for cell-cycle analysis using the ModFit LT^TM^ software package.

### Culture and RNA-sequencing of hPSC differentiation track

Utilizing a modified dual-SMAD inhibition differentiation protocol developed by the Studer laboratory at the Memorial Sloan Kettering Cancer Center, we performed *in vitro* differentiations of hPSCs into SAPs. Over the course of a 40-day differentiation, cells were cultured and sorted on day 16 for the CD49d maker (SOX10 positive cells), when cells are committed to trunc neural crest cells. Cells were harvested at the neural crest and hSAP stages. RNA was isolated from the collected cell pellets by lysing the cells in TRIzol Reagent (ThermoFisher catalog # 15596018) and inducing phase separation with chloroform. Subsequently, RNA was precipitated with isopropanol and linear acrylamide and washed with 75% ethanol. The samples were resuspended in RNase-free water. After RiboGreen quantification and quality control by Agilent BioAnalyzer, 534-850ng of total RNA with DV200% varying from 38-74% was used for ribosomal depletion and library preparation using the TruSeq Stranded Total RNA LT Kit (Illumina catalog # RS-122-1202) according to manufacturer’s instructions with 8 cycles of PCR. Samples were barcoded and run on a HiSeq 4000 in a 50bp/50bp paired end run, using the HiSeq 3000/4000 SBS Kit (Illumina). On average, 48 million paired reads were generated per sample and 35% of the data mapped to the transcriptome.

### RNA-sequencing of perturbated NB cells

Poly-adenylated stranded mRNA sequencing was performed as previously described^13^. In brief, the samples were prepared using the TruSeq Stranded mRNA Sample Prep Kit from Illumina and subsequently sequenced on the Nextseq 500 platform. Sample and read quality was checked with FastQC (v0.11.3). Preprocessing of the fastq reads was performed as previously described^13^. Reads were aligned to the human genome GRCh38 with the STAR aligner (v2.5.3a) and gene count values were obtained with RSEM (v1.2.31). Genes were only retained if they were expressed at counts per million (cpm) above 1 in at least four samples. Counts were normalized with the TMM method (R-package edgeR), followed by voom transformation and differential expression analysis using limma (R-package limma). A general linear model was built with the treatment groups (knockdown or overexpression) and the replicates as a batch effect. Statistical testing was done using the empirical Bayes quasi-likelihood F-test.

Gene Set Enrichment Analysis was performed on the genes ordered according to differential expression statistic value (t). Signature scores were conducted using a rank-scoring algorithm^40^. A custom-made ReplotGSEA function was used to generate gene set enrichment plots. For the data generated on the foetal adrenal glands and differentiation along the sympatho-adrenal lineage, normalisation was done using DESeq2 and rlog transformation, which is more robust in the case when the size factors vary widely.

### Western blot analysis and antibodies

Protein isolation and western blot was performed as previously described^13^. The membranes were probed with the following primary antibodies: anti-SOX11 antibody (SOX11-PAb, 1:1000 dilution), anti-c-MYB antibody (12319S, Cell Signaling, 1:1000 dilution), anti-MYCN antibody (SC-53993, Santa Cruz 1:1,000 dilution), anti-SMARCC1 antibody (11956S, Cell Signaling 1:1,000), and anti-SMARCA4 antibody (3508S, Cell Signaling, 1:500). As secondary antibody, we used HRP-labeled anti-rabbit (7074S, Cell Signalling, 1:10,000 dilution) and anti-mouse (7076P2, Cell Signalling, 1:10,000 dilution) antibodies. Antibodies against Vinculin (V9131; Sigma-Aldrich, 1:10,000 dilution), alpha-Tubulin (T5168, Sigma-Aldrich, 1:10,000 dilution) or beta-actin (A2228, Sigma-Aldrich, 1:10,000 dilution) were used as loading control. The rabbit polyclonal antibody, SOX11-PAb, was custom made (Absea biotechnology, China) against the immunogenic peptide p-SOX11C-term DDDDDDDDDELQLQIKQEPDEEDEEPPHQQLLQPPGQQPSQLLRRYNVAKVPASP TLSSSAESPEGASLYDEVRAGATSGAGGGSRLYYSFKNITKQHPPPLAQPALSPASSRSVSTSSS and used for western blot and chromatin immunoprecipitation for SOX11. All antibodies were diluted in milk/TBST (5 % non-fat dry milk in TBS with 0.1 % Tween-20).

### Chromatin immunoprecipitation (ChIP) assay and ATAC-seq

Chromatin immunoprecipitation (ChIP) and Assay for Transposase-Accessible Chromatin (ATAC) using sequencing was performed as previously described^13^. Briefly, for ChIP-seq a total of 10×10^7^ cells were crosslinked with 1% formaldehyde, quenched with 125 mM glycine, lysed and sonicated with the S2 Covaris for 30 min to obtain 200-300 bp long fragments. Chromatin fragments were immunoprecipitated overnight using 1 *μ*g antibody of SOX11-PAb antibody and 20 *μ*l Protein A UltraLink^®^ Resin (Thermo Scientific) beads per 10×10^6^ cells. Reverse crosslinking was done at 65°C for 15h and chromatin was resuspended in TE-buffer, incubated with RNase A and proteinase K. DNA was isolated and concentration was measured using the Qubit^®^ dsDNA HS Assay Kit. Library prep was done using the NEBNExt Ultra DNA library Prep Kit for Illumina (E7370S) with 500 ng starting material and using 8 PCR cycles according to the manufacturer’s instructions. For ATAC-seq, 50,000 cells were lysed and fragmented using digitonin and Tn5 transposase. The transposed DNA fragments were amplified and purified using Agencourt AMPure XP beads (Beckman Coulter). ChIP-seq and ATAC-seq libraries were sequenced on the NextSeq 500 platform (Illumina) using the Nextseq 500 High Output kit V2 75 or 150 cycles (Illumina).

### ChIP-seq and ATAC-seq data-processing and analysis

Prior to mapping to the human reference genome (GRCh37/hg19) with bowtie2, quality of the raw sequencing data of both ChIP-seq and ATAC-seq was evaluated using FastQC and adapter trimming was done using TrimGalore. Quality of aligned reads were filtered using min MAPQ 30 and reads with known low sequencing confidence were removed using Encode Blacklist regions. Peak calling was performed using MACS2 taking a q value of 0.05 as threshold and peaks were filtered for chr2p amplified regions in the case of IMR-32 cells. DiffBind was used for differential ATAC-peak analysis. Homer^41^ was used to perform motif enrichment analysis, with 200 bp around the peak summit as input. Overlap of peaks, annotation, heatmaps and pathway enrichment was analysed using DeepTools, the R package ChIPpeakAnno, and the web tool enrichR. Sushiplot was used for visualization of the data upon RPKM normalization or log likelihood ratio calculation with MACS2.

## DATA availability

The RNA-sequencing, ChIP-sequencing and ATAC-sequencing datasets generated during this study were deposited in the ArrayExpress database at EMBL-EBI (www.ebi.ac.uk/arrayexpress) with accession numbers: E-MTAB-9338, E-MTAB-9340, E-MTAB-9463 and E-MTAB-9464. Tumor data that supports the findings of this study are available from the Neuroblastoma Research Consortium (NRC, GSE85047), Su et al. [GSE45547]^34^ and Depuydt et al. [GSE103123]^9^, and ChIP-seq data are available from E-MTAB-6562 and E-MTAB-6570.

## ACKNOWLEDGMENTS

The authors would like to thank C. Nunes, L. Mus, K. Verboom, S. Claeys, J. Van Laere and E. De Smet for technical assistance, Lucia Lorenzi for providing the data of the RNA atlas, and P. Reynolds and M. Hogarty for providing the COG-N-373 cell line. We acknowledge the use of the Integrated Genomics Operation Core, funded by the NCI Cancer Center Support Grant (CCSG, P30 CA08748), Cycle for Survival, and the Marie-Josée and Henry R. Kravis Center for Molecular Oncology. This research was supported by the following funding agencies: the Belgian Foundation against Cancer (project 2015-146 and F/2018/1246) to F.S., Ghent University (BOF10/GOA/019 and BOF16/GOA/23) to F.S., the Belgian Program of Interuniversity Poles of Attraction (IUAP Phase VII-P7/03) to F.S., the Fund for Scientific Research Flanders (Research projects G053012N, G050712N and G051516N to F.S., G021415N to K.D and F.S.), ‘Kom op tegen Kanker’ (Stand up to Cancer) the Flemish cancer society (Research grant to F.S.), the European Union H2020 (OPTIMIZE-NB GOD9415N and TRANSCAN-ON THE TRAC GOD8815N to F.S.) and FP7 (ENCCA 261474 and ASSET 259348 to F.S.), ‘Kinderkankerfonds’ (Research grant to F.S.), Olivia Fund to F.S. and Villa Joep to F.S. The following authors B.D., A.L., S.V. and C.V. are supported by an FWO grant.

## AUTHOR CONTRIBUTIONS

B.D., A.L., S.L., S.VHo., S.D., W.V., G.D., C.V., E.S., and N.R. contributed to the development and design of methodology; B.D., C.V., E.D., J.R., J.V., D.C. and J.K. performed computational and statistical analysis; B.D., A.L., S.L., S.D., F.D., S.VHa. and E.S. performed experiments; R.V., J.N., J.K., N.R., M.F., J.S. and S.E. provided material, data and analysis tools, B.D. managed the maintenance of data, B.D., A.L., F.S. and K.D. wrote the original draft, S.L., S.VHo., C.V., J.N., J.K., W.V., S.S.R., T.P. and P.V. contributed to manuscript review and editing, B.D., A.L. and G.D. contributed to data representation and visualization; F.S. and K.D. directed the project and were responsible for funding.

## Supplementary Tables

**Supplementary Table 1:** Differentially expressed genes (adj.P.Val < 0.05) upon *SOX11* knockdown in IMR-32 cells or *SOX11* overexpression in SH-EP cells

**Supplementary Table 2:** SOX11 and MYCN ChIP-seq targets in IMR-32 cells (MACS2, q.Val < 0.05, gene annotation with homer), homer motif enrichment (known motifs) 200 bp size around peak summit, for enhancer or promoter binding

**Supplementary Table 3:** 88-gene signature of SOX11 top direct targets (up upon SOX11 overexpression, down upon SOX11 knockdown, ChIP-seq target and positively correlated expression in 2 NB tumor cohorts)

**Supplementary Table 4:** Cell lines used in the manuscript with the sample ID, origin and MYCN amplification status.

## Supplementary Information

**Supplementary Figure 1:**
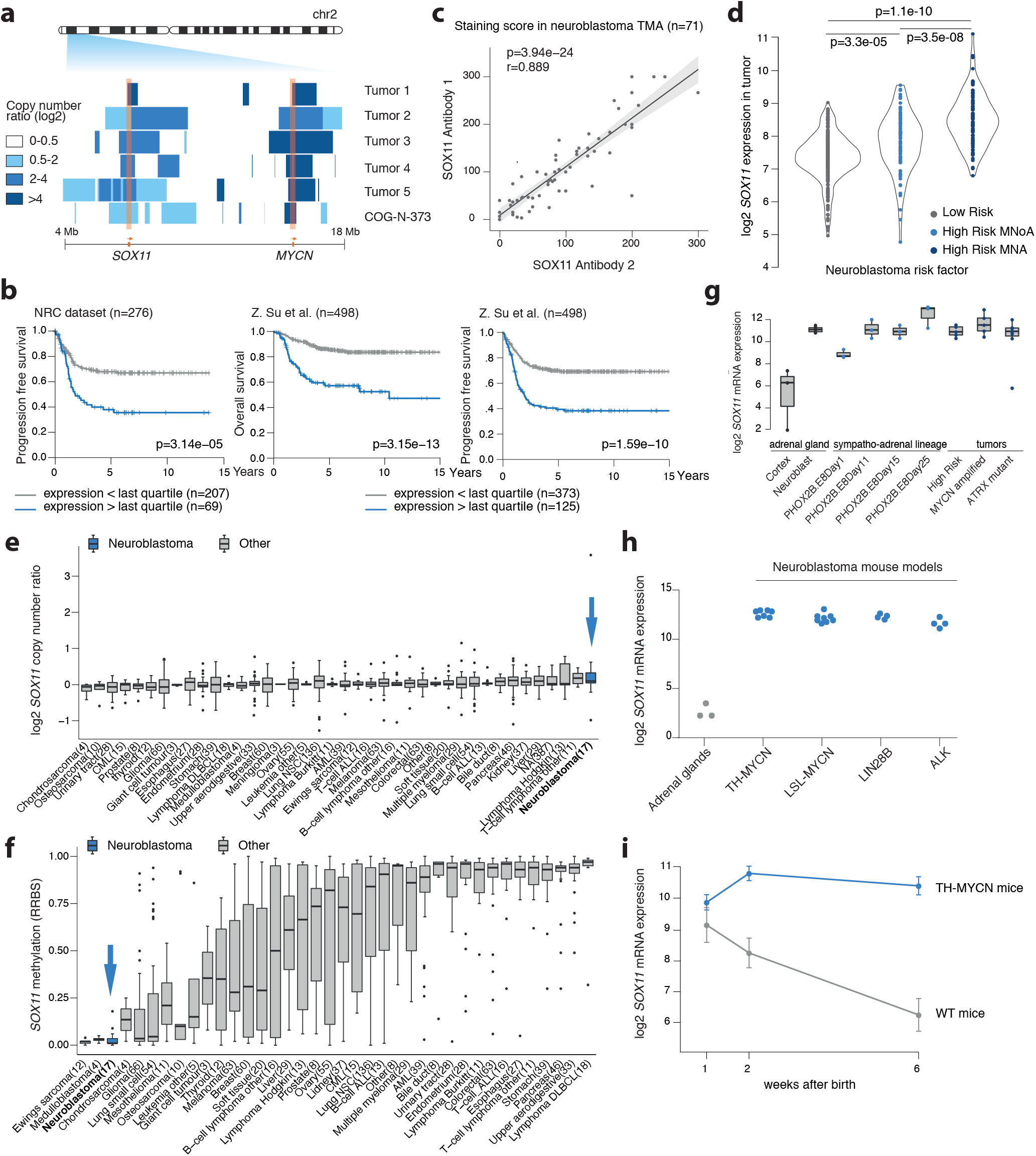
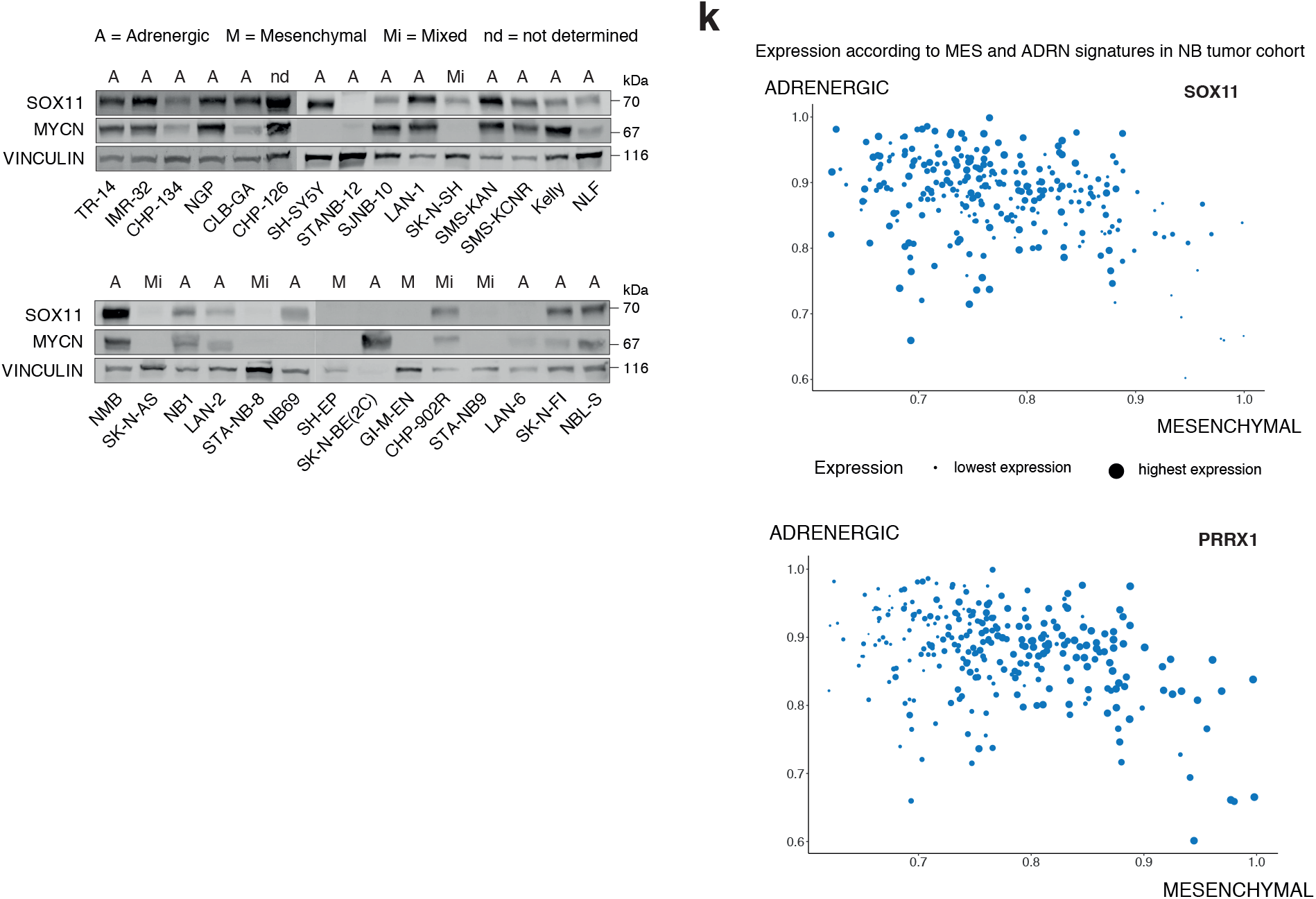
Rare focal amplifications and lineage-specific overexpression of *SOX11* in NB. **a.** Log2 copy number ratio on chr2p (2-18 Mb, hg19), encompassing *SOX11* and *MYCN*, for the 5 NB tumors and 1 NB cell line depicted in Fig 1A. **b.** Kaplan-Meier analysis of 276 (GSE85047) and 498 neuroblastoma patients (GSE62564) with high or low *SOX11* expression (highest quartile cut-off). **c.** IHC for SOX11 with two independent SOX11 antibodies on a TMA of NB tumors showing correlation of SOX11 levels (n=71, r=889, p=3.94e-24, Spearman correlation) confirming specificity. **d.** S*OX11* (log2) expression in *MYCN* amplified high risk tumors as compared to *MYCN* non-amplified high risk tumors and low risk tumors (GSE62546, n=498). ANOVA test followed by Tukey post-hoc test. **e**. Copy number ratio (log2) of *SOX11* in pan-cancer dataset (CCLE) with high copy number for *SOX11* in NB (blue) as compared to other tumor entities (gray). **f.** Genome-wide methylation profile evaluated by reduced representation bisulfite sequencing (RRBS) of *SOX11* in CCLE dataset. *SOX11* show a low methylation profile in NB (blue) as compared to other entities (gray). **g.** *SOX11* (log2) expression in neuroblasts compared to cortex tissue in 3 foetal adrenal glands, during induced differentiation of human sympatho-adrenal precursor cells along the sympatho-adrenal lineage and in NB tumors. **h.** *SOX11* (log2) expression in tumors of *ALK*, *LIN28B*, *LSL-MYCN* and *TH-MYCN* mouse models and mouse adrenal glands. **i.** *SOX11* (log2) mRNA expression in sympathetic ganglia containing hyperplastic lesions and advanced tumors of TH-MYCN^+/+^ mice during MYCN driven NB tumor formation (lightblue) and in normal sympathetic ganglia (darkblue). Error bars represent standard deviation of 4 biological replicates (moderated t-test of Limma Voom, adjusted P value = 7.63e-6). **j.** MYCN, SOX11 and Vinculin protein levels in 29 NB cell lines, annotated as MES-type (M), MIXED-type (Mi), ADRN-type (A) or not dermined (nd) according to Van Groningen et al.^2^, Boeva et al.^3^, and the ADRN/MES signature on mRNA expression level (Fig 2D). **k.** Activity score (rankSum) for ADRN and MES signature^2^ in 283 NB tumors (NRC NB tumor cohort, GSE85047) based on mRNA expression levels. *SOX11* (adrenergic, top) and *PRRX1* (mesenchymal, bottom) expression is represented for each tumor by size. Boxplots in Figure D, E and I show the 1st quartile up to the 3rd quartile of the data and median is a line within the box. Whiskers show the outer two quartiles maximized at 1.5 times the size of the box.

**Supplementary Figure 2:**
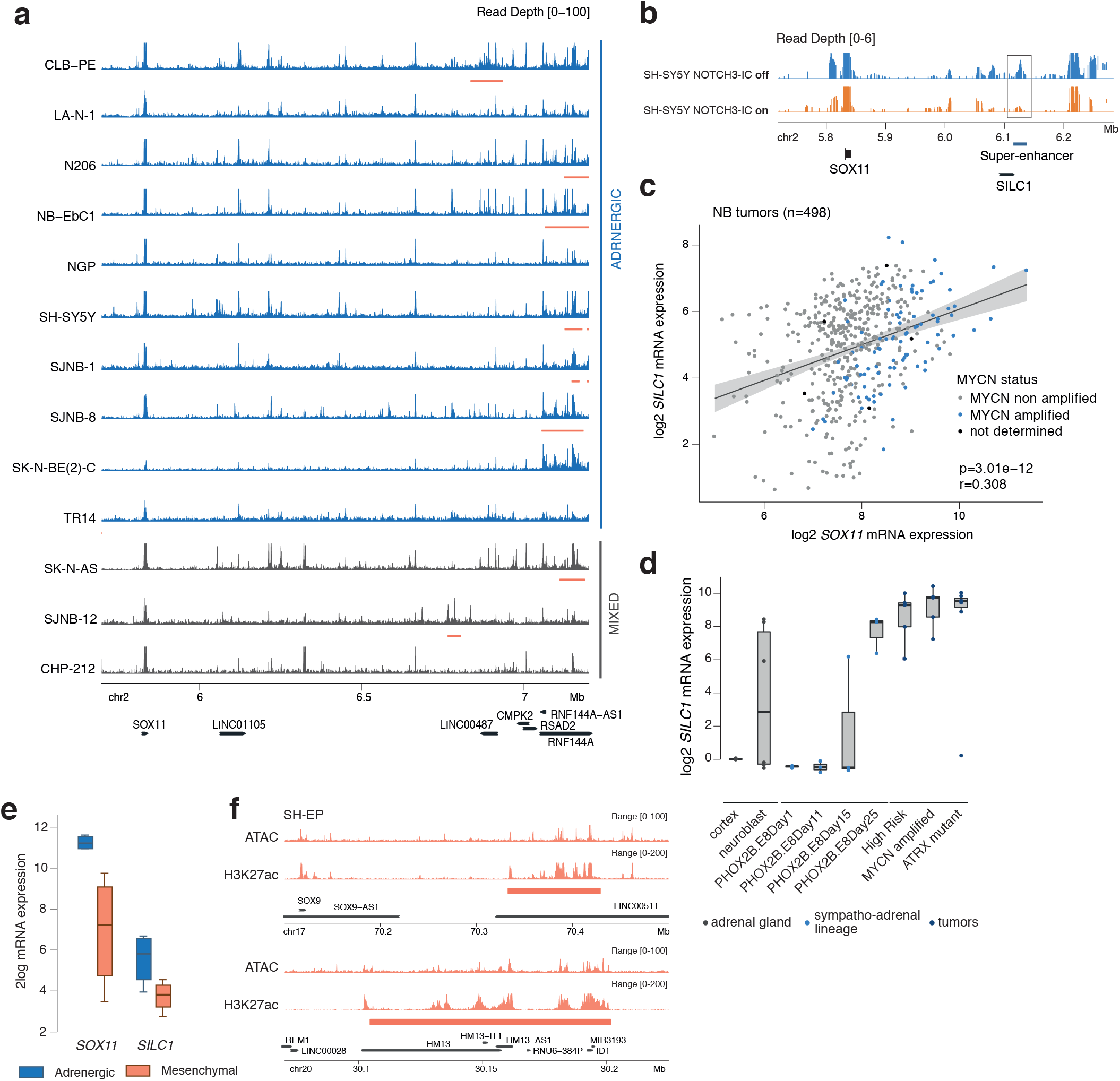
The SOX11 locus is flanked by multiple adrenergic specific enhancers. **a.** H3K27ac activity in a region downstream of SOX11 (chr2, 5.7–7.0Mb, hg19) in 11 adrenergic (NB-EBc1, NGP, CLB-PE, CHP-212, LAN-1, N-206, SH-SY5Y, SJ-NB1, SJ-NB8, SK-N-BE(2c), TR-14) and 1 mixed (SK-N-AS) cell line (blue). Signal represents RPKM normalised ChIP signal, super-enhancers are annotated using LILY (red bar). **b.** H3K27ac activity in a region downstream of SOX11 (chr2, 5.73-6.28Mb, hg19) in SH-Y5Y cells with or without NOTCH3-IC overexpression, the latter resulting in transition from ADRN-type to MES-type cells. The position of the SOX1 downstream super-enhancer is annotated with a rectangle. **c.** SOX11 (log2) mRNA expression correlated with SILC1 log2 mRNA levels in primary NB tumors (n=497, p=3.01e-12, r=0.308, Spearman correlation). MYCN status is depicted for each tumor. **d.** SILC1 (log2) expression in neuroblasts as compared to cortex tissue in 3 fetal adrenal glands, during induced differentiation of human sympatho-adrenal precursor cells along the sympatho-adrenal lineage and in NB tumors **e.** SOX11 and SILC1 log2 expression in the adrenergic subtypes (blue) derived from the isogenic pairs compared to the mesenchymal subtypes (orange) (n=4 pairs). **f.** ATAC-seq and H3K27ac ChIP-seq in SHEP-cells for the super-enhancer regulated mesenchymal markers SOX9 and ID1, showing the validity of the SHEP H3K27ac ChIP-seq and ATAC-seq data used in figure 2E. Signal represents RPKM normalised ChIP signal, super-enhancers are annotated using LILY (red bar). For figure D and E, boxplots are drawn as a box, containing the 1st quartile up to the 3rd quartile of the data values. The median is represented as a line within the box. Whiskers represent the values of the outer two quartiles maximized at 1.5 times the size of the box.

**Supplementary Figure 4:**
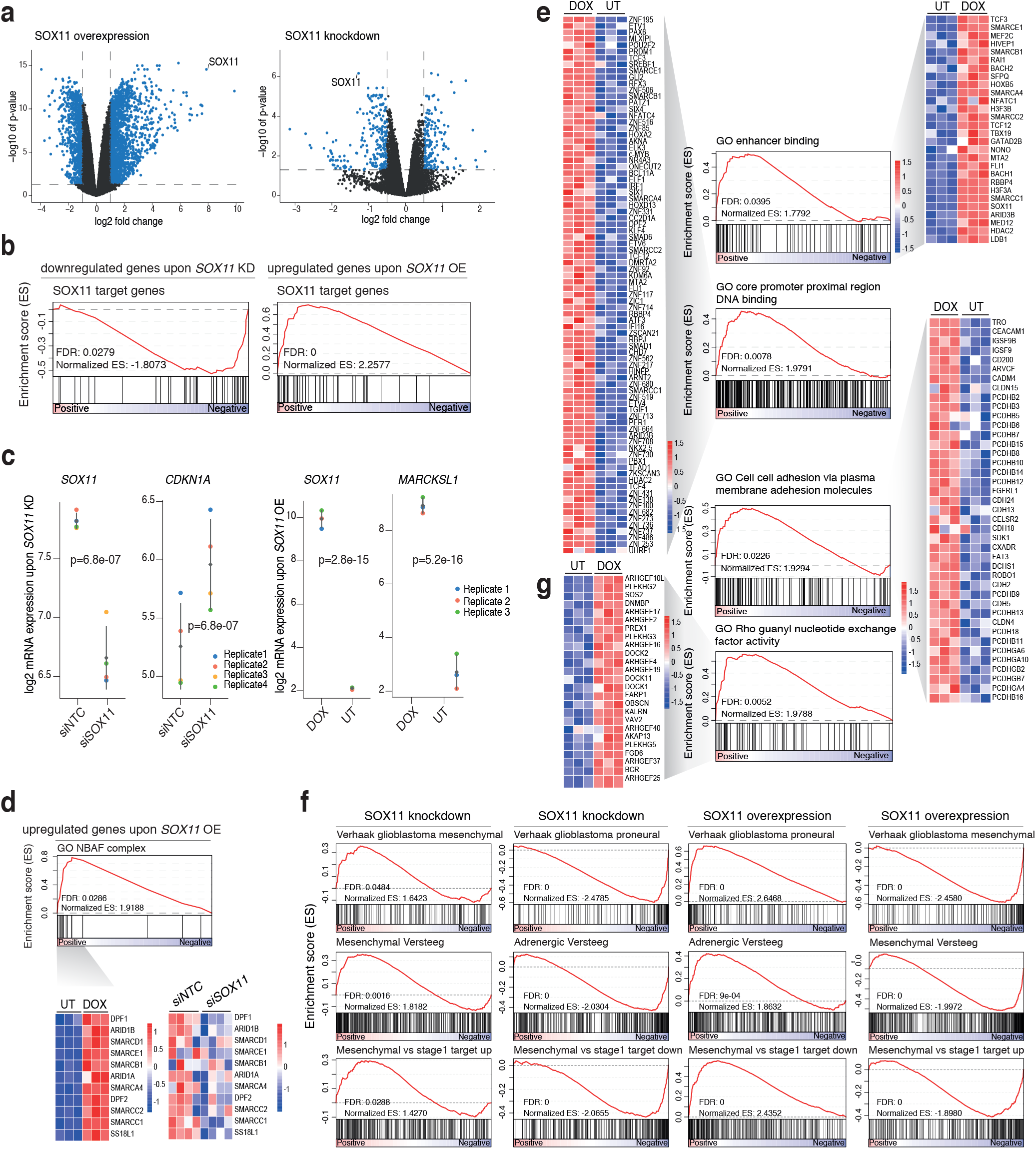
Cell motility, epigenetic control and translation are the hallmarks of the SOX11 regulated transcriptome. **a.** Volcanoplot representing the genes differentially expressed upon *SOX11* overexpression in SH-EP (left, blue, adj p value < 0.05, log FC > 1 or < −1) and *SOX11* knockdown in IMR-32 (right, blue, adj p value < 0.05, log FC > 0.5 or < −0.5). **b**. The MsigDB geneset SOX11 target genes (with genes containing one or more binding sites in their promoter) is highly enriched in the downregulated genes upon *SOX11* knockdown in IMR-32 cells and in the upregulated genes upon *SOX11* overexpression in SH-EP cells, which is validating the RNA-seq data. **c.** *SOX11* and *CDKN1A* log2 mRNA expression levels upon *SOX11* knockdown in IMR-32 cells and *SOX11* and *MARCKSL1* log2 mRNA expression levels upon *SOX11* overexpression in SH-EP cells. Error bars and diamond shape represent respectively the standard deviation and mean of the four biological replicates. Statistical analysis with moderated t-test of Limma Voom (si*NTC* vs si*SOX11*: *SOX11* p=6.8e-07, *CDKN1A* p=6.8e-07; DOX vs UT: *SOX11* p=2.8e-15, *MARCKSL1* p=5.2e-16). **d.** Enrichment of the GO geneset "GO nBAF complex" (neuron-specific BAF or SWI/SNF complex) in the upregulated genes upon *SOX11* overexpression in SH-EP cells. The expression of the representing leading edge is depicted in heatmaps upon *SOX11* overexpression in SH-EP (DOX treated vs Untreated) as well as upon *SOX11* knockdown in IMR-32 cells (siSOX11 vs siNTC). **e.** Enrichment of the GO genesets "GO enhancer binding' and " GO core promoter proximal region DNA binding" in the upregulated genes upon *SOX11* overexpression in SH-EP cells. The expression of the representing leading edges depicted in heatmaps. **f.** Enrichment of the Proneural and Mesenchymal genesets in glioblastoma^18^, the Adrenergic and Mesenchymal genesets in neuroblastoma^2^ and the mesenchymal vs NB stage 1^42^ genesets in the down- and up-regulated genes upon *SOX11* knockdown in IMR-32 and SOX11 overexpression in SH-EP. **g.** Enrichment of the GO genesets "GO Cell cell adhesion via plasma membrane adhesion molecules" and "GO Rho guanyl nucleotide exchange factor activity", in the upregulated genes upon *SOX11* overexpression in SH-EP cells. The expression of the representing leading edges is depicted in heatmaps.

**Supplementary Figure 5:**
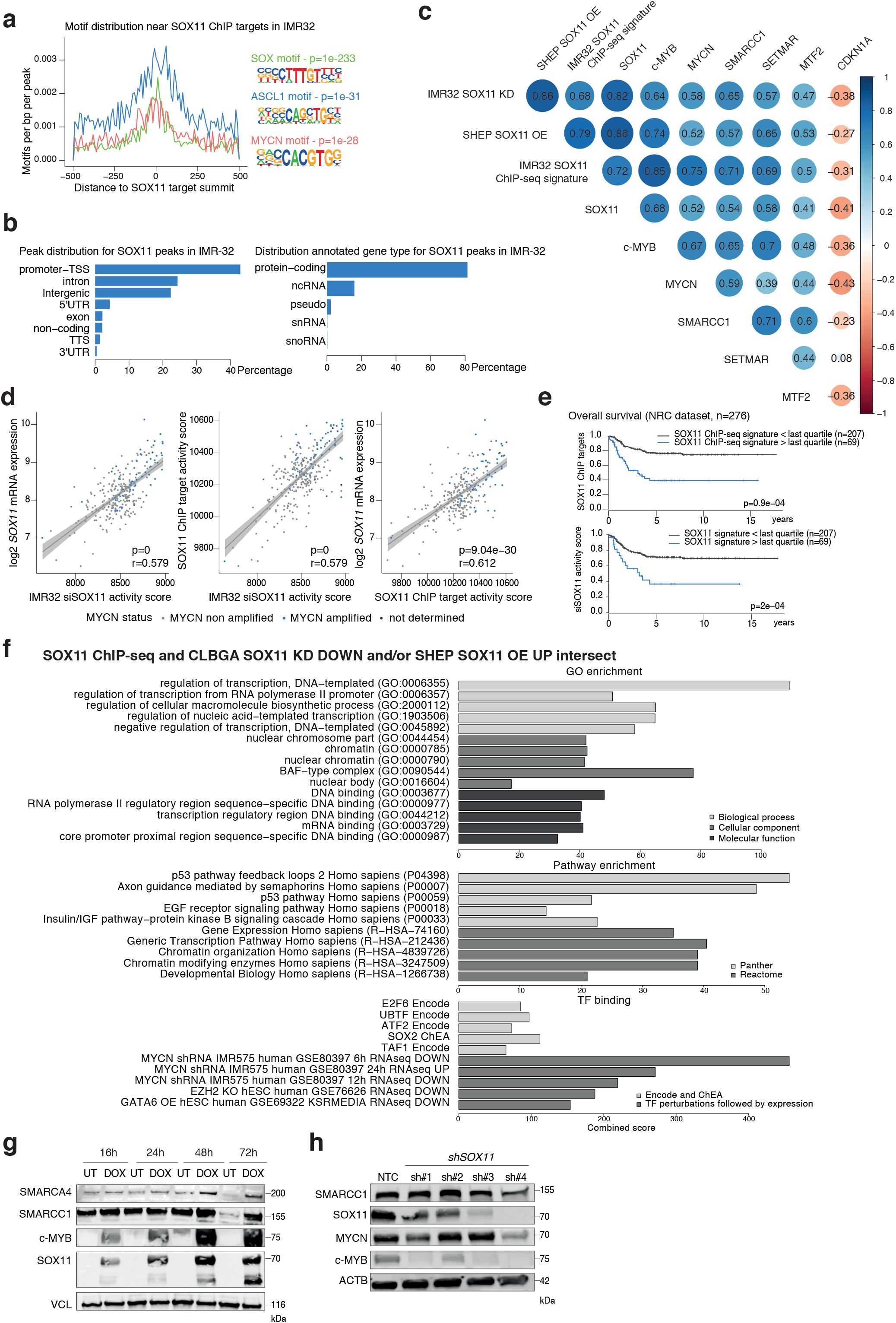
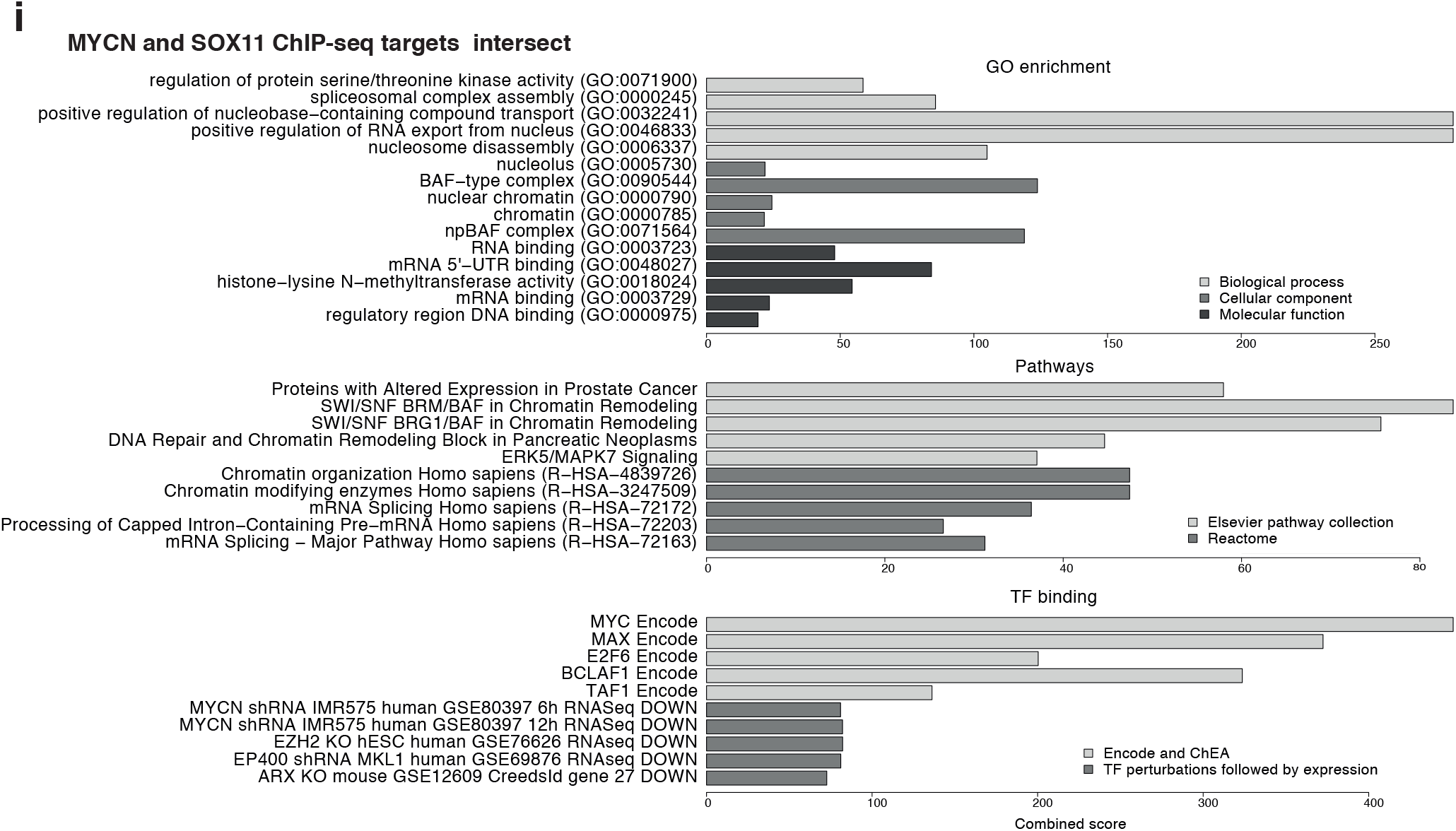
SOX11-mediated chromatin remodeling of lineage specific enhancers. **a.** Distribution of SOX motif, non-canonical E-box motif ASCL1, and canonical E-box motif MYCN at SOX11 peaks in IMR-32 cells (motifs per bp per peak). Signal is depicted −500 to +500 bp from the peak summit. P-value is determined by Homer (SOX motif p=1e-233, ASCL1 motif p=1e-31, MYCN motif p=1e-28). **b.** Genome-wide peak and gene annotation distribution (%) (Homer annotation) for the SOX11 peaks called in IMR-32 cells (MACS2, q<0.05). **c.** Correlation matrix representing the correlation of si*SOX11* signature in IMR-32 and *SOX11* overexpression signature in SH-EP, SOX11 ChIP-seq targets in IMR-32 (all rank sum activity score) and the ranked expression of *SOX11* and affected genes c-*MYB*, *MYCN*, *SETMAR*, *MTF2*, *SMARCC1* and *CDKN1A* in a panel of 29 NB cell lines. The size of dots and color represents the correlation coefficient (Spearman correlation), for the correlations with p<0.05. **d.** Correlation of activity score (rankSum) of the si*SOX11* signature, ChIP target signature in IMR-32 and *SOX11* mRNA expression levels in the NRC NB tumor cohort (*n* = 283, GSE85047) (Left: p=0, r=0.579; middle: p=0, r=0.579; right: p=9.04e-30, r=0.612; Spearman correlation). *MYCN* status is depicted for every tumor sample. For figure C and D, *SOX11* expression was omitted from the si*SOX11* signature score to avoid bias. **e.** Correlation of the activity score (rankSum) of ChIP targets in IMR-32 (top) and the *SOX11* regulated mRNAs signature (bottom) in IMR-32 with overall survival in the NRC NB tumor cohort (*n* = 283, GSE85047). **f.** EnrichR analysis (http://amp.pharm.mssm.edu/Enrichr/) for the SOX11 ChIP targets downregulated after SOX11 knockdown and upregulated after SOX11 overexpression. **g.** SMARCA4, SMARCC1, c-MYB and SOX11 protein levels and loading control Vinculin in an additional clone of SH-EP cells upon *SOX11* overexpression over time. **h.** SOX11, SMARCC1, c-MYB and MYCN protein levels and loading control ACTB in NGP cells upon knockdown of SOX11 using 4 different shRNAs and a non-targeting control (NTC). **i.** EnrichR analysis (http://amp.pharm.mssm.edu/Enrichr/) for the regions overlapping between the SOX11 and MYCN ChIP targets. For F and I, combined Z-score is depicted based on the multiplication of log p value computed with Fisher exact test and the z-score which is the deviation from the expected rank by the Fisher exact test.

**Supplementary Figure 6:**
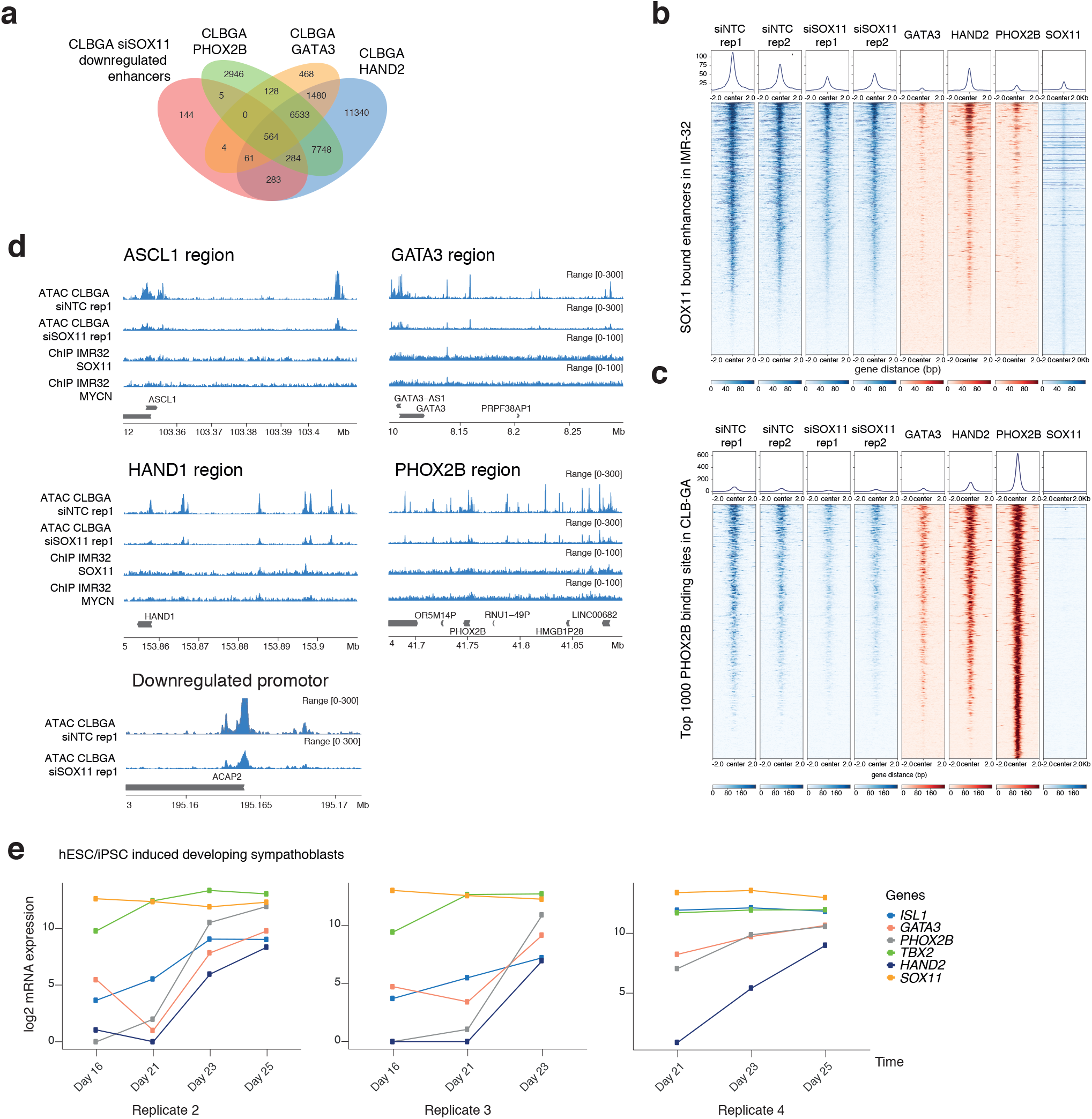
SOX11-mediated chromatin remodeling of lineage specific enhancers. **a.** Overlap (min. overlap = 20 bp) of enhancers with lost chromatin accessibility upon SOX11 knockdown, and PHOX2B, HAND2 and GATA3 binding in the CLB-GA cell line. Binding has only been taking into account when overlapping for at least 50% with H3K27ac and H3K4me1 in parental CLB-GA cells. **b.** Heatmap profiles −2 kb and +2 kb around the summit of SOX11 bound enhancers in IMR-32. The heatmaps represent the RPKM signals for ATAC-seq upon SOX11 knockdown (2 biological replicates), GATA3, HAND2, PHOX2B and SOX11 ChIP-seq profiles, ranked according to the sums of the peak scores across all peaks in the heatmap. Density profiles are shown representing the average ATAC- or ChIP-seq signal at the presented region. **c.** Heatmap profiles 2 kb and +2 kb around the summit of the top 1000 PHOX2B binding sites in CLB-GA. The heatmaps represent the RPKM signals for ATAC-seq upon SOX11 knockdown (2 biological replicates), GATA3, HAND2, PHOX2B and SOX11 ChIP-seq profiles, ranked according to the sums of the peak scores across all peaks in the heatmap. Density profiles are shown representing the average ATAC- or ChIP-seq signal at the presented motifs. **d.** ATAC-seq profiles upon *SOX11* knockdown and control cells, as well as ChIP-seq profiles for SOX11 and MYCN at super-enhancer regions in adrenergic cells (RPKM normalised). **e.** ISL1, GATA3, PHOX2B, TBX2, HAND2 and SOX11 (log2) expression during induced differentiation of hPSC cells along the sympatho-adrenal lineage. Expression levels depicted starting from day 16, when cells are sorted by SOX10 expression levels indicating cells committed to truncal neural crest cells, and followed-up during SAP development stadia until day 25, for three additional replicates (differentiation tracks)

## Notes

### Competing Interest Statement

The authors have declared no competing interest.

## REFERENCES

1 Matthay KK, Maris JM, Schleiermacher G, Nakagawara A, Mackall CL, Diller L et al. Neuroblastoma. Nat Rev Dis Primers 2016; 2: 16078.

2 van Groningen T, Koster J, Valentijn LJ, Zwijnenburg DA, Akogul N, Hasselt NE et al. Neuroblastoma is composed of two super-enhancer-associated differentiation states. Nat Genet 2017; 49: 1261–1266.

3 Boeva V, Louis-Brennetot C, Peltier A, Durand S, Pierre-Eugène C, Raynal V et al. Heterogeneity of neuroblastoma cell identity defined by transcriptional circuitries. Nat Genet 2017; 49: 1408–1413.

4 Garraway LA, Sellers WR. Lineage dependency and lineage-survival oncogenes in human cancer. Nat Rev Cancer 2006; 6: 593–602.

5 Wegner M. From head to toes: the multiple facets of Sox proteins. Nucleic Acids Res 1999; 27: 1409–1420.

6 Potzner MR, Tsarovina K, Binder E, Penzo-Méndez A, Lefebvre V, Rohrer H et al. Sequential requirement of Sox4 and Sox11 during development of the sympathetic nervous system. Development 2010; 137: 775–784.

7 Bhattaram P, Penzo-Méndez A, Sock E, Colmenares C, Kaneko KJ, Vassilev A et al. Organogenesis relies on SoxC transcription factors for the survival of neural and mesenchymal progenitors. Nat Commun 2010; 1: 9.

8 Dodonova SO, Zhu F, Dienemann C, Taipale J, Cramer P. Nucleosome-bound SOX2 and SOX11 structures elucidate pioneer factor function. Nature 2020; 580: 669–672.

9 Depuydt P, Boeva V, Hocking TD, Cannoodt R, Ambros IM, Ambros PF et al. Genomic Amplifications and Distal 6q Loss: Novel Markers for Poor Survival in High-risk Neuroblastoma Patients. J Natl Cancer Inst 2018; 110: 1084–1093.

10 Kumps C, Fieuw A, Mestdagh P, Menten B, Lefever S, Pattyn F et al. Focal DNA copy number changes in neuroblastoma target MYCN regulated genes. PLoS ONE 2013; 8: e52321.

11 Molenaar JJ, Domingo-Fernández R, Ebus ME, Lindner S, Koster J, Drabek K et al. LIN28B induces neuroblastoma and enhances MYCN levels via let-7 suppression. Nat Genet 2012; 44: 1199–1206.

12 Hnisz D, Abraham BJ, Lee TI, Lau A, Saint-André V, Sigova AA et al. Super-enhancers in the control of cell identity and disease. Cell 2013; 155: 934–947.

13 Decaesteker B, Denecker G, Van Neste C, Dolman EM, Van Loocke W, Gartlgruber M et al. TBX2 is a neuroblastoma core regulatory circuitry component enhancing MYCN/FOXM1 reactivation of DREAM targets. Nat Commun 2018; 9: 4866.

14 van Groningen T, Akogul N, Westerhout EM, Chan A, Hasselt NE, Zwijnenburg DA et al. A NOTCH feed-forward loop drives reprogramming from adrenergic to mesenchymal state in neuroblastoma. Nat Commun 2019; 10: 1530.

15 Perry RB-T, Hezroni H, Goldrich MJ, Ulitsky I. Regulation of Neuroregeneration by Long Noncoding RNAs. Mol Cell 2018; 72: 553–567.e5.

16 Durbin AD, Zimmerman MW, Dharia NV, Abraham BJ, Iniguez AB, Weichert-Leahey N et al. Selective gene dependencies in MYCN-amplified neuroblastoma include the core transcriptional regulatory circuitry. Nat Genet 2018; 50: 1240–1246.

17 Meyers RM, Bryan JG, McFarland JM, Weir BA, Sizemore AE, Xu H et al. Computational correction of copy number effect improves specificity of CRISPR-Cas9 essentiality screens in cancer cells. Nat Genet 2017; 49: 1779–1784.

18 Verhaak RGW, Hoadley KA, Purdom E, Wang V, Qi Y, Wilkerson MD et al. Integrated genomic analysis identifies clinically relevant subtypes of glioblastoma characterized by abnormalities in PDGFRA, IDH1, EGFR, and NF1. Cancer Cell 2010; 17: 98–110.

19 El Amri M, Fitzgerald U, Schlosser G. MARCKS and MARCKS-like proteins in development and regeneration. J Biomed Sci 2018; 25: 43.

20 Miao Q, Hill MC, Chen F, Mo Q, Ku AT, Ramos C et al. SOX11 and SOX4 drive the reactivation of an embryonic gene program during murine wound repair. Nature Communications 2019; 10: 4042.

21 Hald ØH, Olsen L, Gallo-Oller G, Elfman LHM, Løkke C, Kogner P et al. Inhibitors of ribosome biogenesis repress the growth of MYCN-amplified neuroblastoma. Oncogene 2018. doi:10.1038/s41388-018-0611-7.

22 Fuglerud BM, Ledsaak M, Rogne M, Eskeland R, Gabrielsen OS. The pioneer factor activity of c-Myb involves recruitment of p300 and induction of histone acetylation followed by acetylation-induced chromatin dissociation. Epigenetics Chromatin 2018; 11: 35.

23 Bögershausen N, Wollnik B. Mutational Landscapes and Phenotypic Spectrum of SWI/SNF-Related Intellectual Disability Disorders. Front Mol Neurosci 2018; 11. doi:10.3389/fnmol.2018.00252.

24 Lei I, Gao X, Sham MH, Wang Z. SWI/SNF protein component BAF250a regulates cardiac progenitor cell differentiation by modulating chromatin accessibility during second heart field development. J Biol Chem 2012; 287: 24255–24262.

25 Jubierre L, Soriano A, Planells-Ferrer L, París-Coderch L, Tenbaum SP, Romero OA et al. BRG1/SMARCA4 is essential for neuroblastoma cell viability through modulation of cell death and survival pathways. Oncogene 2016; 35: 5179–5190.

26 Alver BH, Kim KH, Lu P, Wang X, Manchester HE, Wang W et al. The SWI/SNF chromatin remodelling complex is required for maintenance of lineage specific enhancers. Nat Commun 2017; 8: 14648.

27 Saint-André V, Federation AJ, Lin CY, Abraham BJ, Reddy J, Lee TI et al. Models of human core transcriptional regulatory circuitries. Genome Res 2016; 26: 385–396.

28 Mayran A, Sochodolsky K, Khetchoumian K, Harris J, Gauthier Y, Bemmo A et al. Pioneer and nonpioneer factor cooperation drives lineage specific chromatin opening. Nature Communications 2019; 10: 3807.

29 Yuan X, Seneviratne JA, Du S, Xu Y, Chen Y, Jin Q et al. Single-cell RNA-sequencing of peripheral neuroblastic tumors reveals an aggressive transitional cell state at the junction of an adrenergic-mesenchymal transdifferentiation trajectory. bioRxiv 2020;: 2020.05.15.097469.

30 Furlan A, Dyachuk V, Kastriti ME, Calvo-Enrique L, Abdo H, Hadjab S et al. Multipotent peripheral glial cells generate neuroendocrine cells of the adrenal medulla. Science 2017; 357. doi:10.1126/science.aal3753.

31 Malysheva V, Mendoza-Parra MA, Blum M, Gronemeyer H. Highly dynamic chromatin interactions drive neurogenesis through gene regulatory networks. bioRxiv 2018. doi:10.1101/303842.

32 Bonev B, Mendelson Cohen N, Szabo Q, Fritsch L, Papadopoulos GL, Lubling Y et al. Multiscale 3D Genome Rewiring during Mouse Neural Development. Cell 2017; 171: 557–572.e24.

33 Ye M, Ma J, Liu B, Liu X, Ma D, Dong K. Linc01105 acts as an oncogene in the development of neuroblastoma. Oncology Reports 2019; 42: 1527–1538.

34 Su Z, Fang H, Hong H, Shi L, Zhang W, Zhang W et al. An investigation of biomarkers derived from legacy microarray data for their utility in the RNA-seq era. Genome Biol 2014; 15: 523.

35 Sante T, Vergult S, Volders P-J, Kloosterman WP, Trooskens G, De Preter K et al. ViVar: a comprehensive platform for the analysis and visualization of structural genomic variation. PLoS ONE 2014; 9: e113800.

36 Deleye L, Dheedene A, De Coninck D, Sante T, Christodoulou C, Heindryckx B et al. Shallow whole genome sequencing is well suited for the detection of chromosomal aberrations in human blastocysts. Fertil Steril 2015; 104: 1276–1285.e1.

37 Van Roy N, Forus A, Myklebost O, Cheng NC, Versteeg R, Speleman F. Identification of two distinct chromosome 12-derived amplification units in neuroblastoma cell line NGP. Cancer Genet Cytogenet 1995; 82: 151–154.

38 De Brouwer S, De Preter K, Kumps C, Zabrocki P, Porcu M, Westerhout EM et al. Meta-analysis of neuroblastomas reveals a skewed ALK mutation spectrum in tumors with MYCN amplification. Clin Cancer Res 2010; 16: 4353–4362.

39 Nordström L, Andréasson U, Jerkeman M, Dictor M, Borrebaeck C, Ek S. Expanded clinical and experimental use of SOX11-using a monoclonal antibody. BMC Cancer 2012; 12: 269.

40 Fredlund E, Ringnér M, Maris JM, Påhlman S. High Myc pathway activity and low stage of neuronal differentiation associate with poor outcome in neuroblastoma. Proc Natl Acad Sci USA 2008; 105: 14094–14099.

41 Heinz S, Benner C, Spann N, Bertolino E, Lin YC, Laslo P et al. Simple combinations of lineage-determining transcription factors prime cis-regulatory elements required for macrophage and B cell identities. Mol Cell 2010; 38: 576–589.

42 Rajbhandari P, Lopez G, Capdevila C, Salvatori B, Yu J, Rodriguez-Barrueco R et al. Cross-Cohort Analysis Identifies a TEAD4-MYCN Positive Feedback Loop as the Core Regulatory Element of High-Risk Neuroblastoma. Cancer Discov 2018; 8: 582–599.

